# Searching for Consistent Brain Network Topologies Across the Garden of (Shortest) Forking Paths

**DOI:** 10.1101/2021.07.13.452257

**Authors:** Andrea I. Luppi, Helena M. Gellersen, Alex R. D. Peattie, Anne E. Manktelow, David K. Menon, Stavros I. Dimitriadis, Emmanuel A. Stamatakis

## Abstract

The functional interactions between regions of the human brain can be viewed as a network, empowering neuroscientists to leverage tools such as graph theory to obtain insight about brain function. However, obtaining a brain network from functional neuroimaging data inevitably involves multiple steps of data manipulation, which can affect the organisation (topology) of the resulting network and its properties. Test-retest reliability is a gold standard for both basic research and clinical use: a suitable data-processing pipeline for brain networks should recover the same network topology across repeated scan sessions of the same individual. Analyzing resting-state functional Magnetic Resonance Imaging (rs-fMRI) recordings from two test-retest studies across short (45 minutes), medium (2-4 weeks) and long term delays (5-16 months), we investigated the reliability of network topologies constructed by applying 576 unique pipelines to the same fMRI data, obtained from considering combinations of atlas type and size, edge definition and thresholding, and use of global signal regression. We adopted the portrait divergence, an information-theoretic criterion to measure differences in network topology across all scales, enabling us to quantify the influence of different pipelines on the overall organisation of the resulting network. Remarkably, our findings reveal that the choice of pipeline plays a fundamental role in determining how reproducible an individual’s brain network topology will be across different scans: there is large and systematic variability across pipelines, such that an inappropriate choice of pipeline can distort the resulting network more than an interval of several months between scans. Across datasets and time-spans, we also identify specific combinations of data-processing steps that consistently yield networks with reproducible topology, enabling us to make recommendations about best practices to ensure high-quality brain networks.

## Introduction

The human brain is a remarkably complex system, comprising a large number of regions interacting over time. To address this challenge, neuroscientists have turned to network science, whereby brain regions can be viewed as nodes in a network, and their interactions - e.g. statistical correlations from functional MRI (Bullmore and Sporns, 2009; Fox and Raichle, 2007; Hutchison et al., 2013) - correspond to the connections between nodes. This powerful approach leverages graph theory to quantify key aspects of network organisation, which can in turn provide insights about brain function and dysfunction (Bullmore and Sporns, 2009; Park and Friston, 2013; Sporns, 2011).

In particular, resting-state functional Magnetic Resonance Imaging (rs-fMRI) is a very popular imaging tool, due to its wide applicability: being task-free, it can be easily administered even to challenging populations, from in-utero babies (Turk et al., 2019) to severely injured and even unconscious patients (Demertzi et al., 2019; Huang et al., 2020; Luppi et al., 2019). Since temporal fluctuations of the blood-oxygen-level dependent (BOLD) signal are organized across spatially discrete brain areas (Biswal et al., 1995; Fox and Raichle, 2007), their functional interactions over time (“functional connectivity”) can be quantified in terms of temporal similarities (Hutchison et al., 2013; Smith et al., 2013), resulting in functional brain networks that tabulate the exhaustive number of functional interactions between every possible pair of brain areas (Bassett and Sporns, 2017; Petersen and Sporns, 2015).

Recent methodological developments in network neuroscience and their application to understanding brain disorders have provided powerful insights about both healthy and pathological cognition and brain organisation (Stam, 2014),(Bassett and Sporns, 2017) (Fornito et al., 2015). Aberrant functional brain patterns and dysfunction of subnetworks have been observed with various approaches in many neurological and psychiatric conditions (Hallquist and Hillary, 2019) such as epilepsy (Burianová et al., 2017), Alzheimer’s disease (Greicius et al., 2004),(Johnson et al., 2013), autism (Holiga et al., 2019), schizophrenia (Karbasforoushan and Woodward, 2012) and disorders of consciousness (Demertzi et al., 2019; Huang et al., 2020; Luppi et al., 2019) among others (Petrella, 2011); (Hallquist and Hillary, 2019).

However, recent studies highlighted how different teams and analysis workflows can come to different conclusions about the same neuroimaging dataset (Botvinik-Nezer et al., 2020), owing to a vast pool of possible methodological choices which effectively constitute a combinatorial explosion problem (Eickhoff et al., 2018). Crucially, such a combinatorial explosion also plagues network analyses: even beyond substantial differences introduced by data preprocessing and denoising procedures (Carp, 2012),(Parkes et al., 2018), a wide variety of approaches have been proposed to derive brain networks from preprocessed fMRI data, encompassing choices about the definition of network nodes and edges, edge selection/thresholding, and whether the network should be binary or weighted (Hallquist and Hillary, 2019) (Korhonen et al., 2021), highlighting the intricacies of this issue. Thus, to ensure the value of graph-based estimates as clinical biomarkers, it is of high priority to establish what is the most appropriate way to construct a functional brain network from rs-fMRI data.

One way to identify suitable data-processing steps for brain network construction is in terms of minimising test-retest variability: since it is known that the individual functional connectome is stable over relatively long periods of time (Shehzad et al., 2009), an appropriate pipeline for brain network analysis should produce similar results across repeated scans of the same individual, without introducing distortions in the resulting network and its properties. Existing scientific work compared different network construction steps, considering aspects such as length of acquisition, but also definitions of network nodes using a specific atlas, different ways of quantifying network edges as functional connections, and various edge-filtering schemes, among others (Andellini et al., 2015; Arslan et al., 2018; Braun et al., 2012; Cao et al., 2019, 2014; Du et al., 2015; Ran et al., 2020; Romero-Garcia et al., 2012; Saad et al., 2012; Termenon et al., 2016; Wang et al., 2011, 2017; Welton et al., 2015). Typically, these previous studies focused on specific global or local network properties (e.g. modularity, small-world character, global or local efficiency, down to individual edges) and evaluated the different network construction steps by maximizing the intra-class correlation (ICC) of the adopted global/local network properties (Andellini et al., 2015; Arslan et al., 2018; Braun et al., 2012; Cao et al., 2019, 2014; Du et al., 2015; Ran et al., 2020; Romero-Garcia et al., 2012; Saad et al., 2012; Termenon et al., 2016; Wang et al., 2011, 2017; Welton et al., 2015).

However, these approaches both have limitations. On the one hand, focusing on local aspects (individual edges, node-level properties) runs the risk of “missing the forest for the trees” (Saad et al., 2012), because networks are more than just collections of edges: rather, the way that edges are organised gives rise to micro-, meso- and macro-scale structure, which is precisely what makes network-based approaches so powerful. On the other hand, focusing on specific high-level properties of the network will inevitably limit the generalisability of results, because a vast and ever-growing array of network properties can be defined and used to obtain insights about brain function (Rubinov and Sporns, 2010),(Rubinov and Sporns, 2011)), but there is no guarantee that recommendations pertaining to one will also apply to others.

In the present study, we considered a more general approach to evaluate the various exhaustive sets of combined network construction steps (pipelines): we focus on the network’s topology, that is, the network’s organisation as a whole. For this purpose, we leverage the advantages of the recently introduced “Portrait divergence” (PDiv) measure of dissimilarity between networks (Bagrow and Bollt, 2019). This approach simultaneously takes into account all scales of organisation within a network, from local structure to motifs to large-scale connectivity patterns. Therefore, it incorporates all aspects of network topology, enabling us to go beyond the use of specific and arbitrarily-chosen graph-theoretical properties.

Thus, here we seek to identify network construction pipelines that minimise the difference between brain network topologies of the same individual across different scan sessions. Specifically, we adopt the PDiv to compare functional brain networks obtained from systematic combinations of different options at each step in the network construction process: definition of nodes/brain regions (based on anatomical features, functional homogeneity, or multimodal features); number of nodes (approximately 100, 200, or 400); four different ways to define network edges (from Pearson correlation or mutual information, each either binary or weighted); and eight different approaches to filter the network’s connections (by selecting specific densities, specific thresholds, or using data-driven methods). We also evaluate the effects of a controversial fMRI preprocessing step, the global signal regression (GSR) (Murphy and Fox, 2017) Fig.1 exemplifies the set of choices across the investigated preprocessing steps that influence the construction of a functional brain network, yielding a total set of 576 pipelines (9*4*8*2). To the best of our knowledge, this is the first time in the literature that all these available preprocessing steps are explored simultaneously, and with a focus on topology as a whole rather than on specific network features. To ensure the generalisability of our results (Woo et al., 2017), we further replicate them across three possible time-spans between scans, ranging from minutes to months, in two independent datasets.

**Fig. 1.**
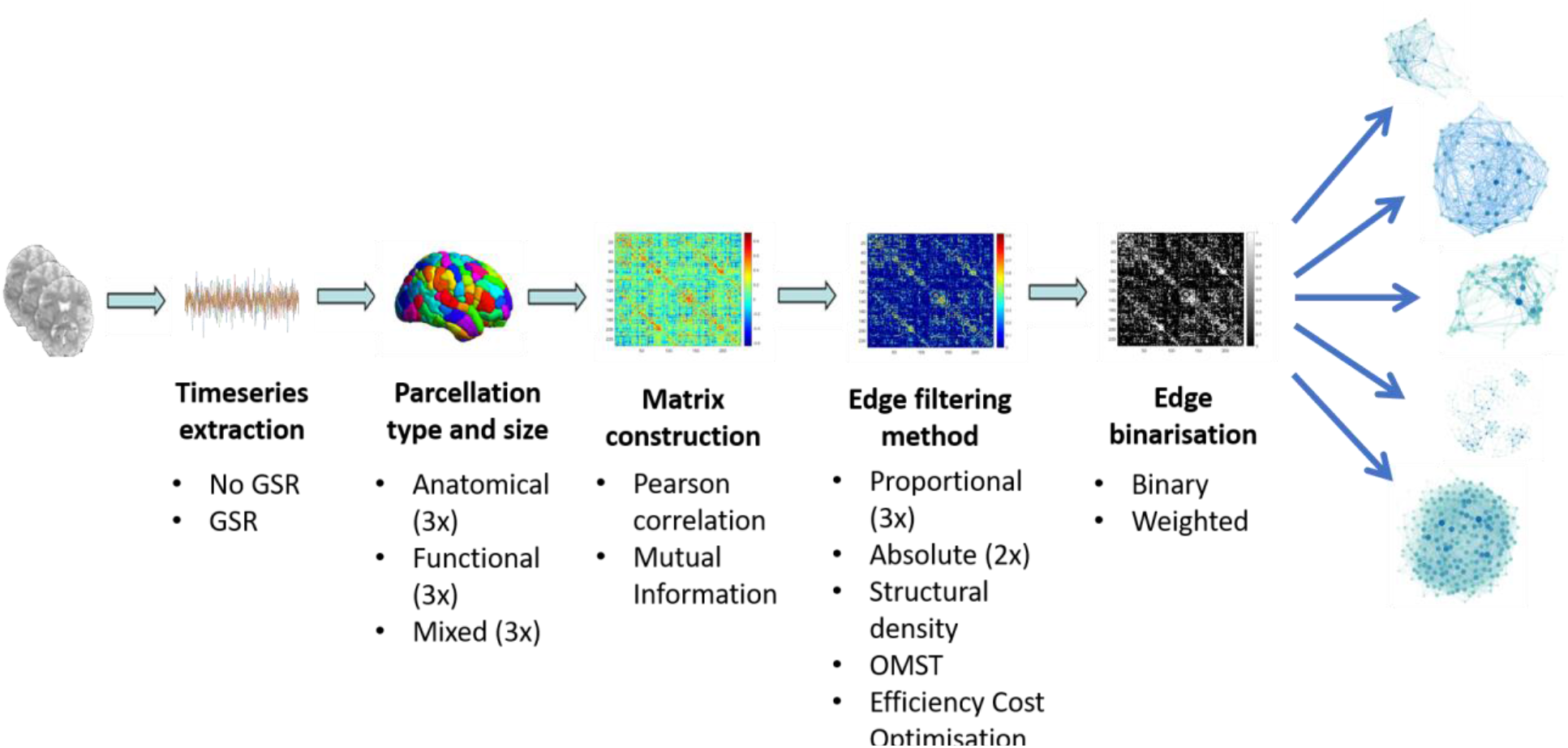
Overview of brain network construction steps. Descriptions outline the alternatives that are considered in the present paper for each step, for the process of obtaining a network from functional MRI data (note that in many cases, this is not an exhaustive sampling of all possibilities that have been presented in the literature).

## Materials and Methods

### NYU Test-Retest dataset

This is an open dataset from the International Neuroimaging Data-Sharing Initiative (INDI) (http://www.nitrc.org/projects/nyu_trt), originally described in (Shehzad et al., 2009). Briefly, this dataset includes 25 participants (mean age 30.7 ± 8.8 years, 16 females) with no history of psychiatric or neurological illness. The study was approved by the institutional review boards of the New York University School of Medicine and New York University, and participants provided written informed consent and were compensated for their participation.

For each participant, 3 resting-state scans were acquired. Scans 2 and 3 were conducted in a single scan session, 45 min apart, which took place on average 11 months (range 5–16 months) after scan 1. Each scan was acquired using a 3T Siemens (Allegra) scanner, and consisted of 197 contiguous EPI functional volumes (TR = 2000 ms; TE = 25 ms; flip angle = 90°; 39 axial slices; field of view (FOV) = 192 × 192 mm2; matrix = 64 × 64; acquisition voxel size = 3 × 3 × 3 mm3). Participants were instructed to relax and remain still with their eyes open during the scan. For spatial normalization and localization, a high-resolution T1-weighted magnetization prepared gradient echo sequence was also obtained (MPRAGE, TR = 2500 ms; TE = 4.35 ms; TI = 900 ms; flip angle = 8°; 176 slices, FOV = 256 mm).

### Cambridge test-retest dataset

Right-handed healthy participants (N=22, age range, 19–57 years; mean age, 35.0 years; SD 11.2; female-to-male ratio, 9/13) were recruited via advertisements in the Cambridge area and were paid for their participation. Cambridgeshire 2 Research Ethics Committee approved the study (LREC 08/H0308/246) and all volunteers gave written informed consent before participating. Exclusion criteria included National Adult Reading Test (NART) <70, Mini Mental State Examination (MMSE) <23, left- handedness, history of drug/alcohol abuse, history of psychiatric or neurological disorders, contraindications for MRI scanning, medication that may affect cognitive performance or prescribed for depression, and any physical handicap that could prevent the completion of testing.

The study consisted of two visits (separated by 2–4 weeks). For each visit, resting-state fMRI was acquired for 5:20 minutes using a Siemens Trio 3T scanner (Erlangen, Germany). Functional imaging data were acquired using an echo-planar imaging (EPI) sequence with parameters TR 2,000 ms, TE 30 ms, Flip Angle 78°, FOV 192 × 192mm2, in-plane resolution 3.0 × 3.0mm, 32 slices 3.0mm thick with a gap of 0.75mm between slices. A 3D high resolution MPRAGE structural image was also acquired, with the following parameters: TR 2,300 ms, TE 2.98 ms, Flip Angle 9°, FOV 256 × 256 mm2. Task-based data were also collected, and have been analysed before to investigate separate experimental questions ( (Vatansever et al., 2015)-(Manktelow et al., 2017). A final set of 18 participants had usable data for both resting-state fMRI scans and were included in the present analysis.

### Functional MRI preprocessing and denoising

Preprocessing of the functional MRI data for both datasets followed the same standard workflow as in our previous studies (Luppi and Stamatakis, 2021), and was implemented in the CONN toolbox (http://www.nitrc.org/projects/conn), version 17f (Whitfield-Gabrieli and Nieto-Castanon, 2012). The following steps were performed: removal of the first 5 volumes to allow for steady-state magnetisation; functional realignment, motion correction, and spatial normalisation to Montreal Neurological Institute (MNI-152) standard space with 2×2×2mm isotropic resolution. Denoising followed the anatomical CompCor method of removing cardiac and motion artifacts, by regressing out of each individual’s functional data the first 5 principal components corresponding to white matter signal, and the first 5 components corresponding to cerebrospinal fluid signal, as well as six subject-specific realignment parameters (three translations and three rotations) and their first- order temporal derivatives (Behzadi et al., 2007). The subject-specific denoised BOLD signal time-series were linearly detrended and band-pass filtered between 0.008 and 0.09 Hz to eliminate both low-frequency drift effects and high-frequency noise.

A further, particularly controversial preprocessing step is global signal regression (GSR): although some authors suggest that GSR may improve subsequent construction of functional brain networks (Braun et al., 2012; Welton et al., 2015), others did not find such an effect (Andellini et al., 2015),(Du et al., 2015) or even reported GSR as deleterious (Cao et al., 2014). Here, we therefore evaluated the effects of including GSR vs omitting it, on subsequent brain network construction.

### Node definition

When deciding how to turn preprocessed and denoised fMRI data into a brain network, the first decision that needs to be made is: what are the elements of the network? Different approaches exist in the literature, including the use of each voxel as a node, to the use of Independent Components Analysis and similar data-driven techniques to obtain study-specific clusterings of brain signals. However, perhaps the most common approach in human network neuroscience is the use of parcellations: pre-defined assignments of neighbouring voxels into regions-of-interest (ROIs). A wide variety of parcellations exist (Eickhoff et al., 2018), and recent work reported how the choice of parcellation scheme can affect aspects such as structure-function similarity estimation (Messé, 2020) but also the intra-subject and inter-subject variability of the whole-brain resting-state modeling (Popovych et al., 2021). Parcellation schemes vary on two main dimensions: the criterion based on which clusters are identified (e.g. based on neuroanatomy, or functional considerations, or a combination thereof) and the number of ROIs - ranging from a few tens to thousands.

Here, we considered both of these dimensions: we employed parcellations spanning three scales (approximately 100, 200 and 400 nodes) and obtained based on anatomical, functional, or multimodal considerations (summarised in Table 1).

**Table 1.** Atlases adopted in the present study, by scale (rows) and method (columns).

|  | Anatomical | Functional | Mixed |
| --- | --- | --- | --- |
| **Scale-100** | Lausanne 129 | Schaefer 100 +<br>Melbourne 16 | AAL 90 |
| **Scale-200** | Lausanne 234 | Schaefer 200 +<br>Melbourne 32 | Brainnetome 246 |
| **Scale-400** | Lausanne 463 | Schaefer 400 +<br>Melbourne 54 | Glasser 360 +<br>Melbourne 54 |

As anatomical parcellations we consider the popular Automated Anatomical Labelling (AAL) atlas, with 90 cortical and subcortical regions (Tzourio-Mazoyer et al., 2002), and the multi-scale Lausanne atlas with 129, 234 and 463 cortical and subcortical nodes obtained by subdividing the sulcus-based Desikan-Killiany atlas (Cammoun et al., 2012).

As functional parcellation we use the recent Schaefer atlas (Schaefer et al., 2018) which combines local gradients and global similarity across task-based and resting-state functional connectivity. Following our previous work, we included versions with 100, 200 and 400 cortical regions, respectively supplemented with 16, 32 or 54 subcortical regions from the recent subcortical functional atlas developed by Tian and colleagues (Tian et al., 2020).

Finally, we included the Brainnetome atlas, which comprises 210 cortical and 36 subcortical regions, identified by combining anatomical, functional and meta-analytic information (Fan et al., 2016) and the Glasser parcellation comprising 360 cortical regions identified by combining multi-modal information about cortical architecture, function, connectivity, and topography (Glasser et al., 2016). The Glasser parcellation was also supplemented with the 54-region version of the Melbourne atlas, in order to include a comparable number of subcortical regions, resulting in 414 ROIs.

### Functional Connectivity

For each parcellation, the average denoised BOLD timeseries across all voxels belonging to a given ROI were extracted. We considered two alternative ways of quantifying the interactions between regional BOLD signal timeseries. First, we used Pearson correlation, whereby for each pair of nodes *i* and *j*, their functional connectivity *F_ij_* was given by the Pearson correlation coefficient between the timecourses of *i* and *j*, over the full scanning length. Second, we also used the mutual information *I*, which quantifies the interdependence between two random variables X and Y, and is defined as the average reduction in uncertainty about X when Y is given (or vice versa, since this quantity is symmetric):

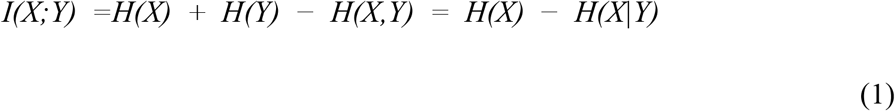

With *H(X)* being the Shannon entropy of a variable X. Unlike Pearson correlation, mutual information considers both linear and nonlinear relationships, and it does not provide negative values. Following previous work, the values in each individual matrix of mutual information were divided by the maximum value in the matrix, thereby normalising them to lie between zero and unity.

### Filtering Schemes

Both Pearson correlation and MI provide continuous values for the statistical association between pairs of nodes, resulting in a dense matrix of functional connections. Therefore, some form of filtering is typically employed to remove spurious connections that are likely to be driven by noise, and obtain a sparse network of functional connectivity. However, there is no gold standard approach to decide which connections to retain, and different filtering schemes have emerged in the literature. Here, we considered 8 different edge filtering schemes (Table 2), described below.

**Table 2.**
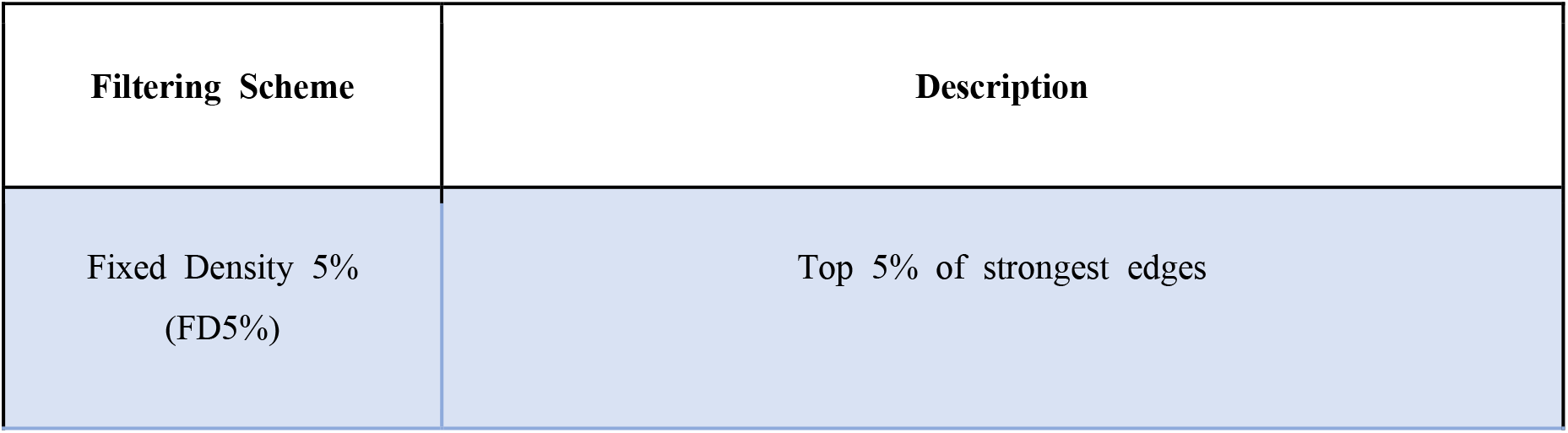
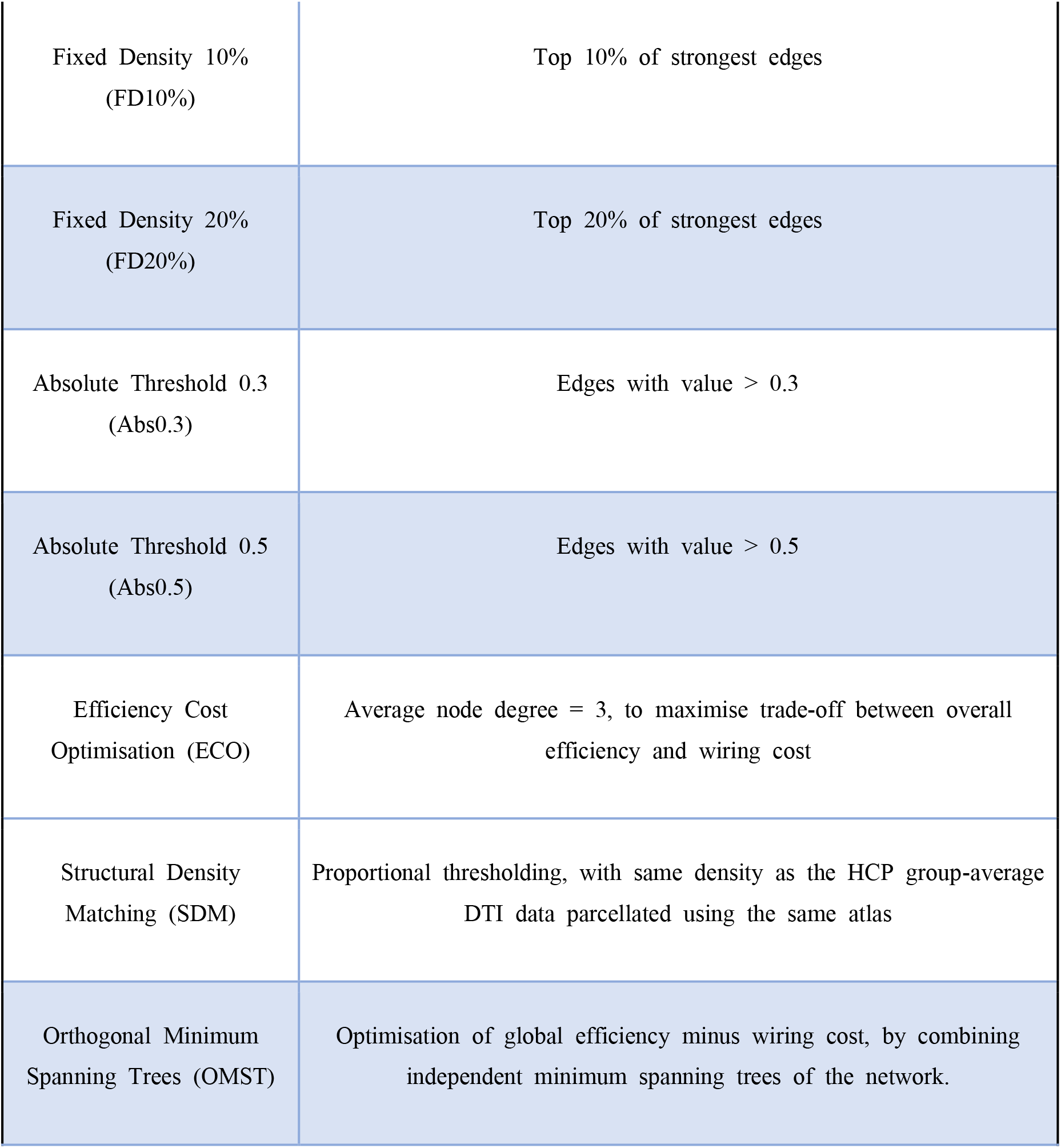
Edge filtering schemes adopted in the present study.

### Absolute Thresholding

The simplest approach to decide which edges to retain is to accept or reject edges based on a pre-determined minimum acceptable weight. However, there is no consensus in the literature about which threshold one should adopt. Here, we considered absolute threshold values of 0.3 or 0.5 (for Pearson correlation, only positively-value edges were considered).

### Proportional Thresholding

Absolute thresholding can produce networks with very different densities, which can introduce confounds in subsequent network analyses. Therefore, a popular approach simply retains a fixed proportion of the strongest edges. However, there is once again no consensus in the literature on the correct proportion of edges to retain. We therefore employed four different density levels, in the range commonly reported in the literature: fixed density (FD) of 5%, 10%, and 20% of the strongest edges.

### Structural Density Matching

The main problem with proportional thresholding is the selection of an appropriate target density - especially since this may vary depending on the number of nodes in the network. To address this issue in a principled manner, we recently introduced a method termed Structural Density Matching (SDM), whereby the proportion of functional edges to retain corresponds to the density *s* of the corresponding structural connectome (the network of anatomical connectivity obtained from the group-averaged diffusion-weighted MRI data from the Human Connectome Project (Yeh et al., 2018). In other words, SDM ensures that functional and structural networks obtained using the same parcellation have the same density, instead of using an arbitrary target density.

### Efficiency Cost Optimisation

The Efficiency Cost Optimisation (ECO) is designed to optimise the trade-off between the network’s overall efficiency (sum of global and average local efficiency) and its wiring cost (number of edges) (De Vico Fallani et al., 2017), by ensuring that the network maximises the following target function *J*:

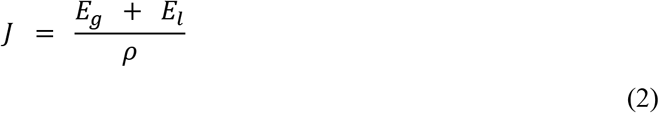

With *E_g_* and *E_l_* being the global and mean local efficiency of the network, respectively. This filtering scheme produces sparse graphs while still preserving their structure, as demonstrated by its empirical success at discriminating between different graph topologies (De Vico Fallani et al., 2017). Here, we obtained ECO-thresholded graphs by setting a proportional threshold such that the average node degree would be 3, since previous analytic and empirical results indicate that the optimal density corresponds to enforcing an average node degree approximately equal to 3 (De Vico Fallani et al., 2017).

### Orthogonal Minimum Spanning Trees

OMST (Dimitriadis et al., 2017a, 2017c) is another data-driven approach intended to optimise the balance between efficiency and density of the network, while also ensuring that the network is fully connected. Specifically, the method involves three steps: (1) identifying the minimum set of edges such that each node can be reached from each other node - known as the minimum spanning tree (MST); (2) identifying an alternative (orthogonal) MST, and combining it with the previous one; (3) repeating steps (1) and (2) until the network formed by the progressive addition of orthogonal MSTs optimises a global cost function defined as *E*_*g*_ – Cost (with Cost corresponding to the ratio of the total weight of the selected edges, divided by the total strength of the original fully weighted graph). This approach produces plausibly sparse networks without imposing an a-priori level across all subjects, and it has been shown that the resulting networks provide higher recognition accuracy and reliability than many alternative filtering schemes (Dimitriadis et al., 2017c).

### Binarisation

For all filtering schemes considered here, edges that were not selected were set to zero. However, edges that were included in the network could be weighted or unweighted. In the case of unweighted (binary) networks, we set all non-zero edges to unity. Otherise, their original weight was retained.

### Topological Distance as Portrait Divergence

To quantify the difference between network topologies, we used the recently developed Portrait Divergence. The Portrait Divergence between two graphs G1 and G2 is the Jensen-Shannon divergence between their “network portraits”, which encode the distribution of shortest paths of the two networks (Bagrow and Bollt, 2019). Specifically, the network portrait is a matrix *B* whose entry *B_lk_, l* = 0, 1, …, *d* (with *d* being the graph diameter), *k* = 0, 1, …, *N* − 1, is the number of nodes having *k* nodes at shortest-path distance *l*.

Thus, to compute the Portrait Divergence one needs to compute the probability *P(k, l)* (and similarly *Q(k, l)* for the second graph) of randomly choosing two nodes at distance *l* and, for one of the two nodes, to have *k* nodes at distance *l*:

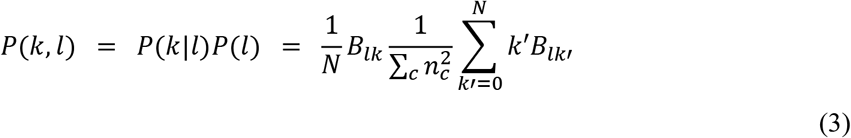

where *n_c_* is the number of nodes in the connected component *c*. Then, the Portrait Divergence distance is defined using the Jensen-Shannon divergence (an information-theoretic notion of distance):

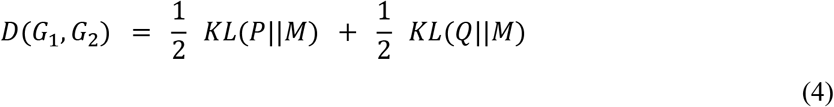

where *M* = *(P + Q)/2* is the mixture distribution of *P* and *Q*, and *KL*(⋅||⋅) is the Kullback-Leibler divergence.

The Portrait Divergence offers three key advantages. First, it is based on network portraits, which do not change depending on how a graph is represented. Comparing network topologies based on such “graph invariants” is highly desirable, because it removes the potential confound of encoding format. Second, the Portrait Divergence does not require the networks in question to have the same number of nodes or edges, and it can be applied to both binary and weighted networks - making it ideally suited for the applications of the present study. And finally, the Portrait Divergence is not predicated on a single specific network property, but rather it takes into account all scales of structure within networks, from local structure to motifs to large scale connectivity patterns: that is, it considers the topology of the network as a whole (Bagrow and Bollt, 2019).

For each subject, at each timepoint, we obtained one brain network following each of the possible combinations of steps above (576 distinct pipelines in total).

For each pipeline, we then computed the Portrait Divergence between networks obtained from the same subject at different points in time, and subsequently obtained a group-average value of Portrait Divergence for each pipeline.

## Results

We used an information-theoretic measure of distance between network topologies across scales, termed Portrait divergence (PDiv), to systematically compare 576 alternative network construction pipelines in terms of their ability to recover similar brain network topologies from functional MRI scans of the same individual across minutes (NYU dataset, same-session scans), weeks (Cambridge dataset), or months (NYU dataset, between-sessions comparison) (see Methods).

Being grounded in information theory, the Portrait divergence between two networks can be interpreted as measuring how much information is lost when using one network to represent another. It ranges from 0 (no information loss) to 1 (complete information loss); in the present case, since the two networks are derived from different scans of the same individual, we aim to identify pipelines that minimise test-retest PDiv.

For each dataset, Fig.2 illustrates the distributions of group-mean test-retest similarities of network topologies (portrait divergence) across the full set of 576 pipelines. Clearly, two patterns can be observed.

**Fig. 2.**
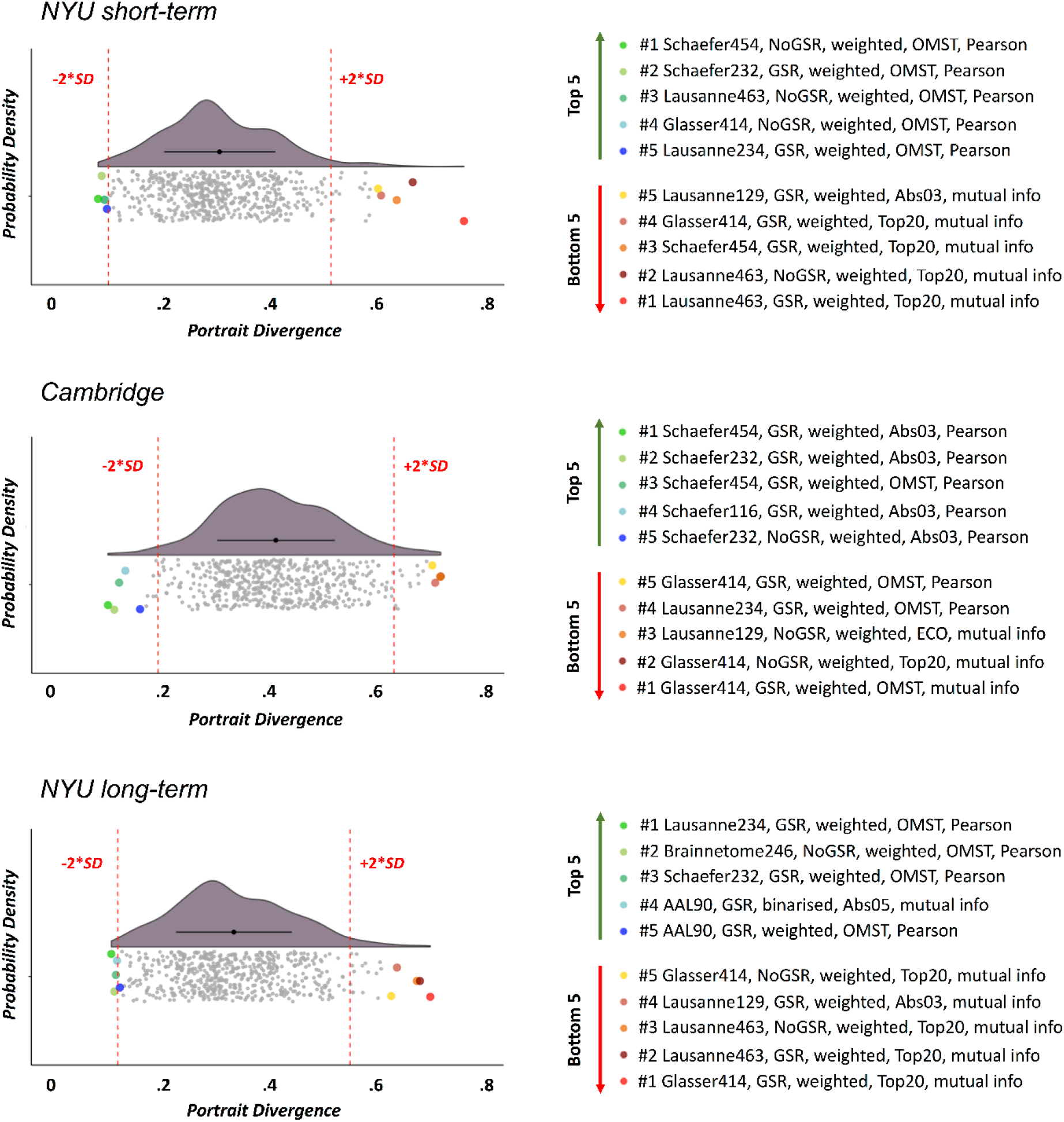
Distribution of group-average portrait divergence values for each of 576 alternative network construction pipelines, across different time intervals. Top: NYU short-term dataset (rescan within 45 minutes). Middle: Cambridge dataset (rescan within 2-4 weeks). Bottom: NYU long-term dataset (rescan within 16 months; average 11.4). Right-side: highlighting the top 5 (lowest PDiv) and bottom 5 performers (highest PDiv). Red lines mark 2 standard deviations from the mean of the distribution. **Abbreviations**. GSR: Global Signal Regression. OMST: Orthogonal Minimal Spanning Trees.

First, network construction pipelines differ widely in how well they are able to recover the same network topology across different scans of the same individual, on average - whether on a timescale of minutes, weeks, or months.

Second, our results indicate substantial consistency across the three time intervals considered here, in terms of which data-processing steps feature prominently among the pipelines that are best (and worst) at minimising the average within-subject PDiv. Additionally, correlation between all pipelines’ ranks across time intervals indicated very high consistency for short-term NYU and long-term NYU (Spearman’s rho = 0.94, p < 0.001). While these two timespans are the most different among those considered here, the data were acquired on the same subjects. However, the pipelines’ ranks obtained from both short- and long-term NYU were also significantly and positively correlated with the ranks obtained in the independent Cambridge dataset (Cambridge vs NYU-short: Spearman’s rho = 0.31, p < 0.001; Cambridge vs NYU-long: Spearman’s rho = 0.32, p < 0.001) (Fig.3), indicating that pipelines’ suitability for network construction is not dataset-specific but rather can generalise to different groups of individuals.

**Fig. 3.**
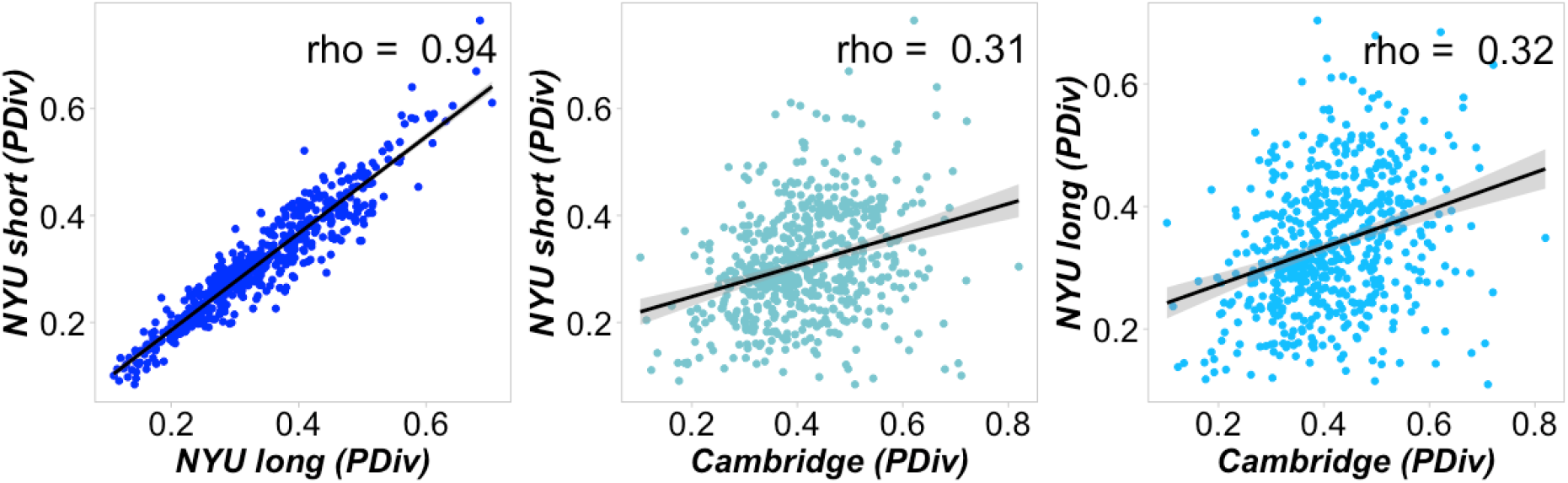
Rank-based correlations of the pipelines’ performance (PDiv) across datasets. Our analysis indicated very high correlations for short-term NYU and long-term NYU, and moderate correlations for the medium-term Cambridge with both long and short NYU). All p < 0.001.

Therefore, it is possible to identify pipelines that perform well regardless of the intervening time-span: Table 3 reports the overall top 20 pipelines, identified as the pipelines with the highest mean rank (lowest test-retest PDiv) across all three datasets. Correspondingly, the overall lowest-ranked pipelines (highest test-retest PDiv) are reported in Table 4. An interactive table with all pipelines and their ranks can be found in the Supplementary Material.

**Table.3.** Overall 20 best-performing pipelines, based on the highest mean rank (lowest test-retest PDiv across the group) across the cohorts.

| Mean Global Rank | Cam Rank | NYUs Rank | NYUI Rank | Global PDiv | Cam PDiv | NYUs PDiv | NYUI PDiv | Atlas Type | Atlas Size | GSR Options | Binarised vs. Weighted | Threshold | Edge Type |
| --- | --- | --- | --- | --- | --- | --- | --- | --- | --- | --- | --- | --- | --- |
| 4 | 7 | 3 | 2 | 0.13 | 0.18 | 0.09 | 0.12 | Functional | Scale 200 | GSR | Weighted | OMST | Pearson |
| 6 | 3 | 9 | 7 | 0.12 | 0.12 | 0.11 | 0.14 | Functional | Scale 400 | GSR | Weighted | OMST | Pearson |
| 10 | 11 | 6 | 12 | 0.15 | 0.19 | 0.12 | 0.13 | Functional | Scale 200 | No GSR | Weighted | OMST | Pearson |
| 12 | 6 | 15 | 16 | 0.15 | 0.17 | 0.12 | 0.15 | Functional | Scale 100 | GSR | Weighted | OMST | Pearson |
| 13 | 4 | 12 | 23 | 0.14 | 0.13 | 0.14 | 0.14 | Functional | Scale 100 | GSR | Weighted | Abs0.3 | Pearson |
| 15 | 10 | 17 | 17 | 0.16 | 0.19 | 0.12 | 0.15 | Anatomical | Scale 200 | No GSR | Weighted | OMST | Pearson |
| 16 | 9 | 24 | 15 | 0.16 | 0.19 | 0.12 | 0.16 | Functional | Scale 100 | GSR | Weighted | OMST | Mutual Info |
| 16 | 19 | 8 | 21 | 0.16 | 0.21 | 0.13 | 0.13 | Functional | Scale 100 | No GSR | Weighted | OMST | Mutual Info |
| 17 | 37 | 5 | 10 | 0.17 | 0.26 | 0.11 | 0.13 | Mixed | Scale 100 | GSR | Weighted | OMST | Pearson |
| 27 | 27 | 22 | 32 | 0.18 | 0.24 | 0.16 | 0.16 | Anatomical | Scale 400 | No GSR | Binarised | FD20% | Mutual Info |
| 31 | 30 | 37 | 26 | 0.19 | 0.24 | 0.15 | 0.18 | Functional | Scale 400 | GSR | Binarised | FD20% | Pearson |
| 32 | 15 | 43 | 39 | 0.18 | 0.21 | 0.16 | 0.18 | Functional | Scale 400 | GSR | Binarised | FD20% | Mutual Info |
| 33 | 74 | 4 | 20 | 0.19 | 0.3 | 0.13 | 0.12 | Mixed | Scale 100 | GSR | Binarised | Abs0.5 | Mutual Info |
| 33 | 51 | 30 | 19 | 0.19 | 0.28 | 0.13 | 0.17 | Functional | Scale 100 | GSR | Weighted | FD20% | Pearson |
| 42 | 60 | 31 | 34 | 0.21 | 0.29 | 0.16 | 0.17 | Mixed | Scale 100 | GSR | Weighted | SDM | Pearson |
| 44 | 18 | 50 | 63 | 0.2 | 0.21 | 0.19 | 0.19 | Functional | Scale 100 | No GSR | Weighted | Abs0.3 | Pearson |
| 47 | 80 | 25 | 35 | 0.21 | 0.3 | 0.16 | 0.16 | Mixed | Scale 100 | No GSR | Weighted | OMST | Pearson |
| 51 | 84 | 38 | 30 | 0.21 | 0.31 | 0.15 | 0.18 | Anatomical | Scale 100 | GSR | Weighted | OMST | Mutual Info |
| 54 | 13 | 79 | 70 | 0.21 | 0.2 | 0.2 | 0.22 | Functional | Scale 400 | GSR | Binarised | Abs0.3 | Pearson |
| 57 | 120 | 39 | 13 | 0.21 | 0.32 | 0.12 | 0.18 | Functional | Scale 100 | GSR | Binarised | Abs0.5 | Mutual Info |

**Abbreviations.** Abs: absolute. ECO: efficiency cost optimisation. FD: fixed density. GSR: global signal regression. OMST: orthogonal minimal spanning trees. SDM: structural density.

**Table.4.**
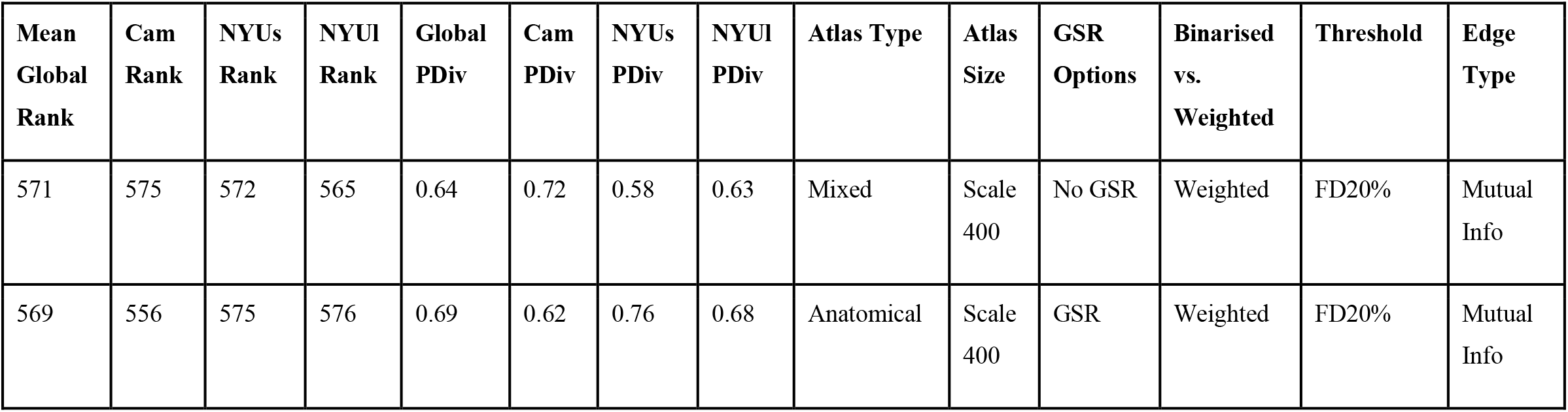
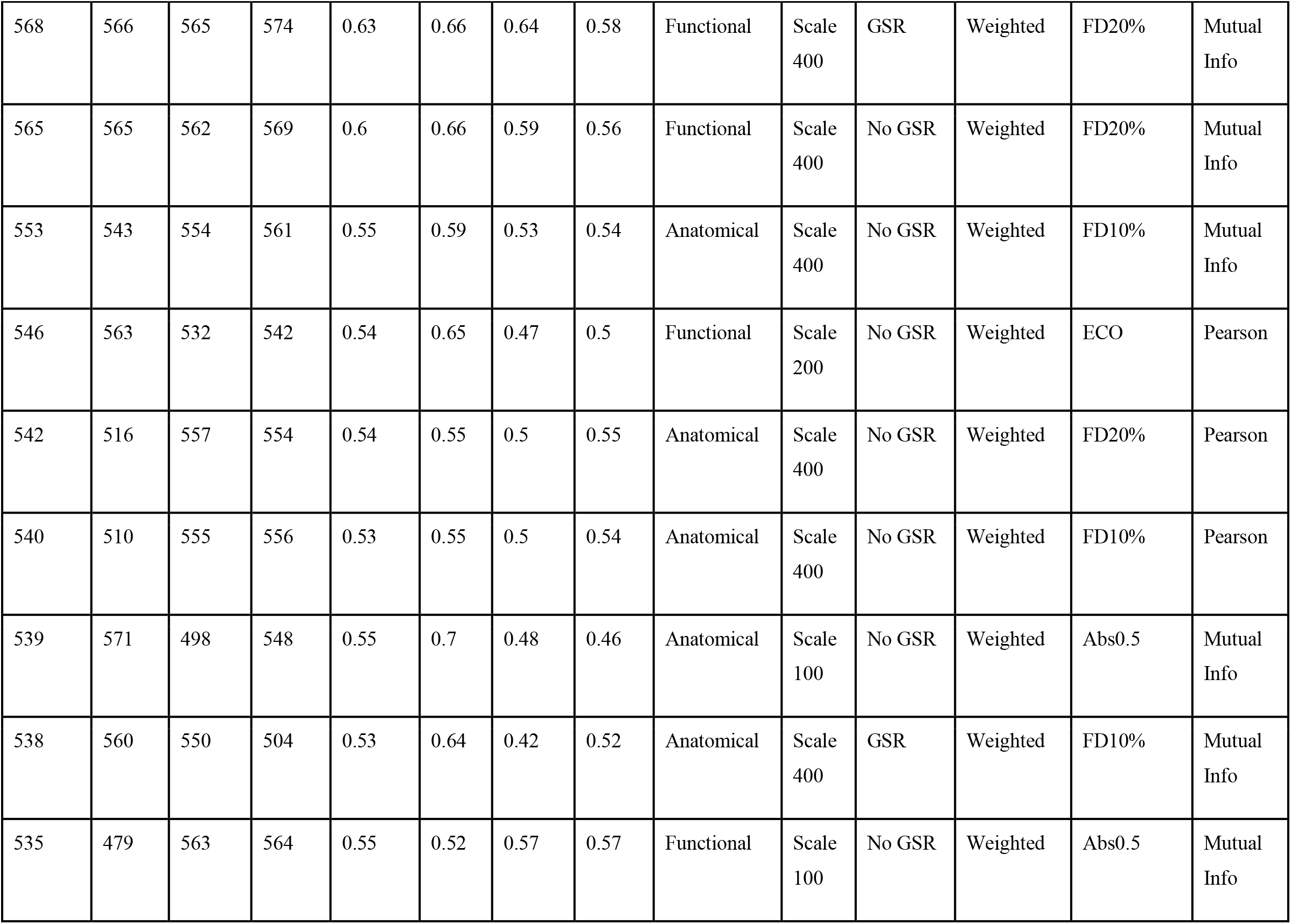
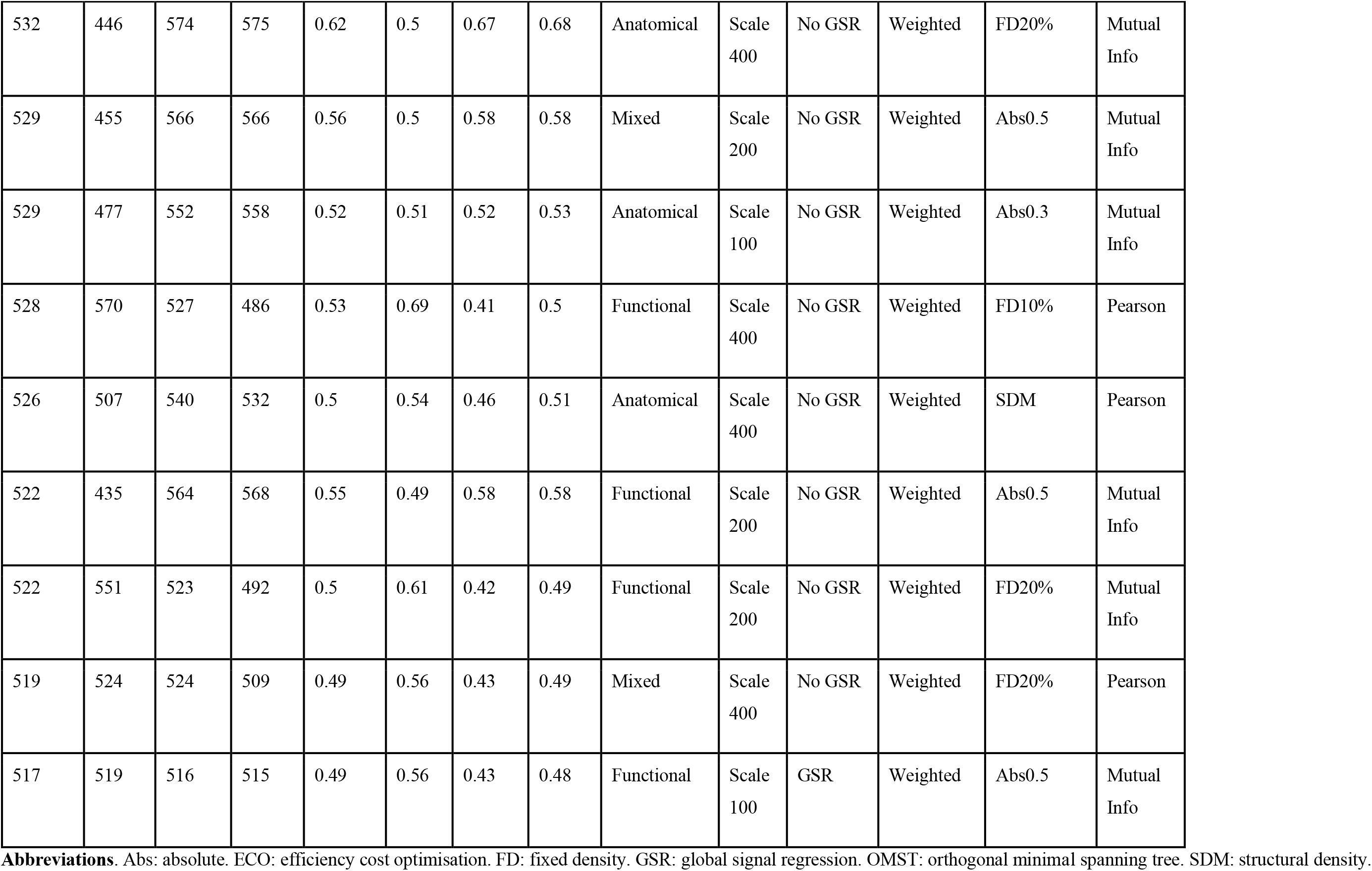
Overall 20 worst-performing pipelines, based on the lowest mean rank (highest test-retest PDiv across the group) across the cohorts.

From this ranking, it becomes clear that whether a pipeline performs well (in terms of minimising the PDIv between different scans of the same individual) is not randomly determined, but rather there are systematic aspects: many of the same individual steps are consistently present across the best-performing pipelines (Fig.4 indicates the relative prevalence of each pipeline option among the 20 overall best- and worst-performing pipelines). Importantly, there is a set of options that consistently feature as part of the pipelines that most faithfully reproduce network topology from different rs-fMRI recordings of the same individual. This set includes:

i. use of global signal regression (GSR) during preprocessing;
ii. node definition based on Schaefer’s functional atlas;
iii. edge definition in terms of Pearson’s correlation coefficient;
iv. the Orthogonal Minimal Spanning Trees (OMST) scheme (Dimitriadis et al., 2017a),(Dimitriadis et al., 2017c) to determine which edges to retain;
v. use of a weighted network.

**Fig. 4.**
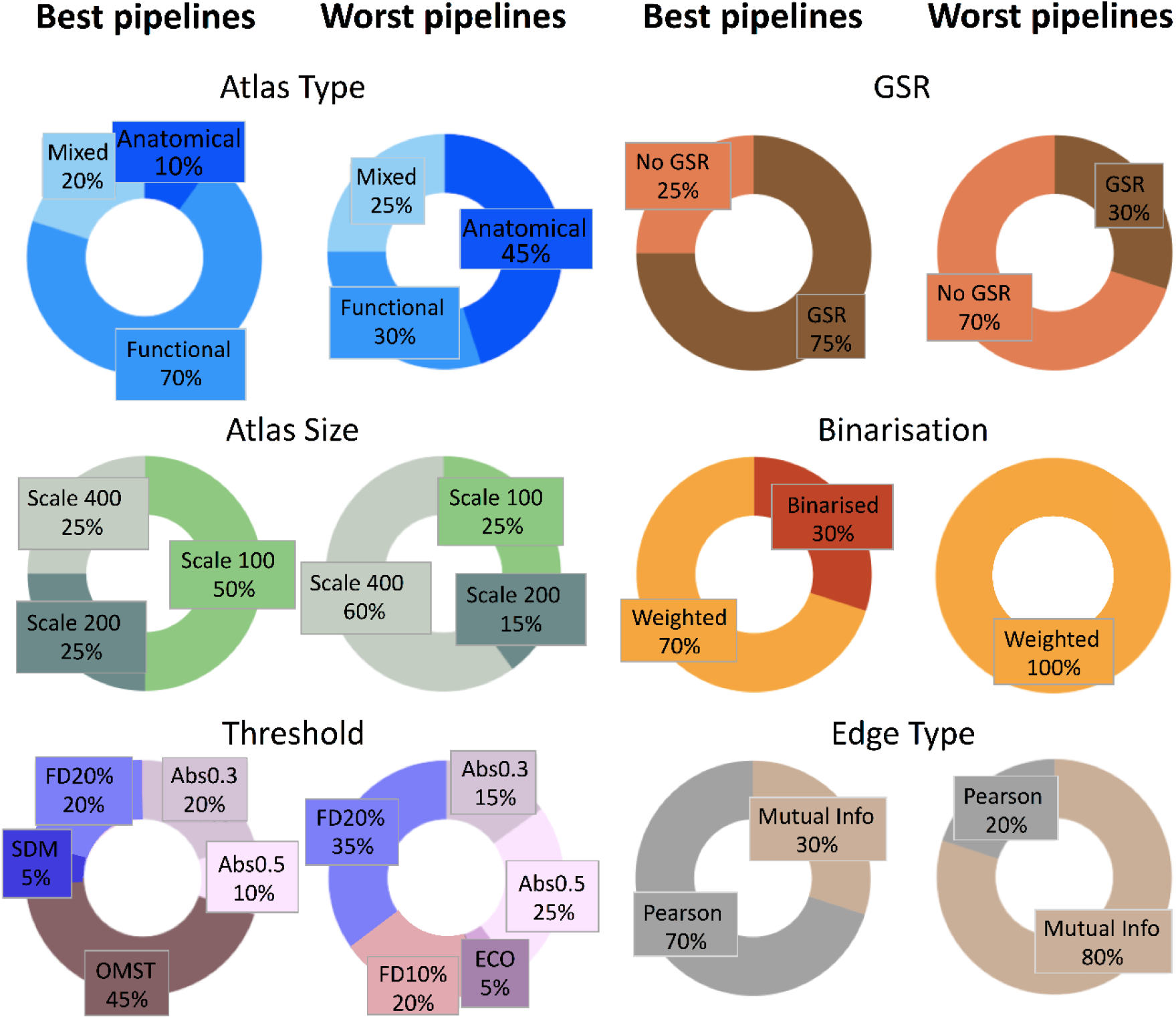
Prevalence of specific network construction steps among the best and worst pipelines. Pie charts demonstrate, for each network construction step, the proportion of each option that is found among the top and bottom 20 pipelines, respectively. **Abbreviations**. Abs: absolute. ECO: efficiency cost optimisation. FD: fixed density. GSR: global signal regression. OMST: orthogonal minimal spanning tree. SDM: structural density.

Consistency was also observed among the worst-performing pipelines across the three time-spans considered (Fig. S3). The set of choices that produces the least reproducible network topologies includes: various atlases, mostly scale-400 parcellations, mutual information connectivity estimator, no GSR and a slight prevalence of 20% fixed-density thresholding criterion for thresholding, without binarisation.

Interestingly, both the best- and worst-performing pipelines involve the use of weighted rather than binary edges. This shows that, when considered in isolation, this step is not indicative of whether the pipeline as a whole will produce good results. Indeed, although the case of binary vs weighted edges is especially obvious, systematically investigating each step revealed that there is large overlap between pipelines that differ in only one step, in terms of their distributions of PDiv values, with consistently large variance (Fig. S1-S6).

Therefore, a pipeline’s performance is not solely attributable to any specific step: rather, it is due to the synergy between combinations of steps. Indeed, in addition to the consistent presence of many specific options among the best pipelines (as shown in Fig.4), there are also systematic patterns in how these options are combined. This is especially evident when focusing on the global top 5 pipelines: all use the Schaefer functional atlas; four out of five use the Schaefer functional atlas with weighted edges determined by Pearson correlation followed by OMST thresholding; of these four, three also use GSR, therefore differing in only one single aspect: the parcellation scale. Therefore, the combination of these steps appears especially recommended for obtaining networks with reliable topology.

Clear systematic combinations of steps are also found among the bottom 5 pipelines globally: all five use a scale-400 parcellation with weighted edges determined from mutual information followed by fixed-density thresholding (out of which, the bottom four use 20% fixed density). Therefore, although no single choice is unequivocally beneficial or deleterious when it is investigated in isolation with respect to the rest of the preprocessing steps, this is no excuse to disregard the issue of which options to adopt: different combinations of options into specific pipelines will have a large impact on the resulting network topology, in a way that is remarkably consistent across datasets and across time-spans.

## Discussion

A tremendous amount of neuroimaging research with functional MRI is devoted to finding reliable brain network-based biomarkers for brain function and its disorders – but this process involves a combinatorial explosion of arbitrary choices (Korhonen et al., 2021). Test-retest reliability of brain network organisation (topology) is a prerequisite to define reliable connectomic biomarkers (Fornito et al., 2015),(Hallquist and Hillary, 2019). Here, we systematically investigated 576 unique pipelines that a neuroscientist could adopt to obtain brain networks from resting-state fMRI data, arising from the combination of several key data-processing steps. Rather than choosing any arbitrary graph-theoretical property for our comparisons, we focused on the networks’ overall topology across all scales, which enabled us to compare the pipelines in terms of their ability to recover similar brain network topologies in data obtained from the same individual across short (minutes), medium (weeks) and long timespans (months), in two independent datasets.

Summarized, our findings reveal large and systematic differences across pipelines in terms of their ability to recover similar topologies for the same individual. These findings were consistent across short, medium and long time-intervals between scan sessions, in two independent datasets. Remarkably, an inappropriate choice of pipeline can greatly impair the ability to recover a reliable network topology: even for scans obtained less than 45 minutes apart, we observed up to a 5-fold increase in topological dissimilarity (PDiv) compared with the best-performing pipelines (Fig. 2). In contrast, the best-performing pipelines will recover similar network topologies for the same individual, even at a distance of several months.

These results suggest that an inappropriate choice of network construction pipeline may have devastating effects for longitudinal studies of brain network properties. More broadly, test-retest reliability of network topology is of fundamental importance for any subsequent graph-theoretical analysis: any pipelines that recover vastly different topologies from two scans of the same individual taken within the same hour, are liable to produce misleading results when used to associate network properties with behavioural traits (Smith et al., 2013) or clinical outcomes such as psychosis, autism or Alzheimer’s disease (Woo et al., 2017). Thus, identification of reliable network construction pipelines represents a fundamental prerequisite for both network-based investigation (Messaritaki et al., 2019; Nichols et al., 2017); (Messaritaki et al., 2019) and subsequent efforts aimed at clinical translation (Chen et al., 2018).

Fortunately, clear patterns emerged across the best-performing pipelines in terms of specific network construction steps,with substantial consistency both in different datasets (NYU and Cambridge) and across different time-spans. Namely, our results lead to the recommendation of global signal regression, Schaefer’s atlas for node definition, Pearson’s correlation as connectivity estimator, weighted edges, and finally Orthogonal Minimum Spanning Trees (OMST) as an appropriate topological filtering scheme.

Reassuringly, many of these recommendations are consistent with the recommendations of previous methodological investigations employing several different approaches. In particular, the use of Schaefer’s functional parcellation for edge definition, and OMST as filtering scheme for weighted networks, both provided topologically representative networks (Luppi and Stamatakis, 2021), in terms of minimising topological differences between pipelines (rather than focusing on topological test-retest reliability between different scans using the same pipeline, as we did here). In other studies, OMST has also outperformed alternative thresholding schemes for functional networks in terms of recognition accuracy and reliability (Dimitriadis et al., 2017a), and more recently also for structural connectivity networks (Dimitriadis et al., 2017b); (Messaritaki et al., 2019)). Likewise, functional atlases have also been recommended over anatomical ones in previous work (Cao et al., 2019), possibly because anatomical atlases can include ROIs with widely different timeseries (Korhonen et al., 2017). Therefore, our results suggest a convergence of recommendations for brain network construction across different criteria - possibly heading towards consistent analytic practices in the field.

In contrast to the best options, a pattern of choices within the worst-performing pipelines include: a large number of nodes (over 400), mutual information as a connectivity estimator with weighted edges, and fixed-density thresholding. Importantly, however, our results do *not* indicate that any of these choices is inappropriate, when considered in isolation. Rather, a key finding of the present study is that no single step uniquely determines a pipeline’s ability (or inability) to accurately recover the network’s topology (Fig S1-S6). Pipelines differing by only one step are largely overlapping in terms of their portrait divergence distributions - to the point that the use of weighted edges features prominently both among the best- and worst-performing pipelines (likely because binarisation removes both noise and neurobiologically relevant information). On the other hand, we see that substantial differences exist at each step in the network construction process. Therefore, it is clear that a pipeline’s suitability for brain network construction is determined in large part by the synergy between its constituent steps, and not just by the choice of one specific step.

Given the importance of considering network construction steps jointly rather than in isolation, it is a key strength of our study that we considered the entire data processing workflow, rather than only focusing on one or two specific steps, as most previous publications did. Another key strength of our approach is that we did not base our investigation on the test-retest reliability of arbitrarily chosen local or global graph-theoretical metrics, which would inevitably limit the generalisability of any results and recommendations. Instead, using the topology-based portrait divergence enabled us to take into account both local and global aspects of network organisation across scales (Bagrow and Bollt, 2019). Therefore, by considering the network’s topology as a whole our results are inherently more general than results based on any specific graph-theoretical metric.

Of course, test-retest reliability is not the only criterion that neuroscientists need to consider for their choice of network construction pipelines: ultimately, the resulting networks need to also demonstrate empirical usefulness by providing neurobiologically meaningful results (Shirer et al., 2015)-(Noble et al., 2019). In particular, although it is important to minimise spurious distortions, it is also important to be able to identify true changes, whether due to neuromodulation (both in terms of tasks and changes in arousal), behavioural history (sleep deprivation, time of day etc), external pharmacology (caffeine, nicotine, prescription or recreational drug use) and clinical characteristics. The ideal pipeline(s) should be sensitive to these differences, while filtering out differences that are merely due to noise. While it is not possible to exclude the presence of such differences in our medium- and long-term retest datasets for individual participants, we expect that such differences should be randomly distributed and thus cancel out at the group level. Additionally, we note that none of these potentially relevant factors are likely to influence functional brain networks obtained with the same hour, as we studied in the short-term NYU dataset; and remarkably, pipelines that minimise topological distance in the short term tend to also do so over longer periods of time. Nevertheless, it will be important for future work to demonstrate the validity of our recommended pipelines, by associating the resulting topology with behavioural or personality traits (Smith et al., 2015) or by increasing the predictive power of a connectomic model to discriminate healthy controls from different brain disorders (Chen et al., 2018).

### Limitations

In this study, we endeavoured to systematically sample and combine many of the most common options across each step in the process of constructing a functional brain network from rs-fMRI data - resulting in 576 unique pipelines. However, due to combinatorial explosion it would be unfeasible to consider every single option that has been proposed in the literature, and this inevitable limitation should be borne in mind when interpreting our results.

In particular, we did not explore potential differences between resting-state conditions (eyes-open vs eyes-closed vs naturalistic viewing) (Van Dijk et al., 2010; Wang et al., 2011), or the impact of scan duration and arousal state (Laumann et al., 2017). Similarly, we did not consider a wide number of alternative parcellation schemes in existence (Arslan et al., 2018; Eickhoff et al., 2018), nor parcellation-free methods for node definition, such as voxelwise (Du et al., 2015), and ICA-based methods (Kiviniemi et al., 2009; Smith et al., 2011). Likewise, many alternative thresholding methods also exist, whether based on statistical significance (Váša et al., 2018), or shrinkage methods (Smith et al., 2011; Wang et al., 2017) or avoiding thresholding entirely, by using analytic methods that can deal with fully connected and signed networks (Rubinov and Sporns, 2011).

It is also known that different motion correction strategies can have an effect on the validity of BOLD signals and subsequent network characteristics; however, no correction strategy offered perfect motion correction (Parkes et al., 2018). Here, we adopted a widely used denoising strategy (aCompCor), and only considered the additional effects of global signal regression due to ongoing controversy about its effect on functional connectivity (Murphy and Fox, 2017; Saad et al., 2012). Our data-driven results suggest a beneficial effect of GSR on topological test-retest reliability, presumably due to the removal of noise confounds. However, investigators are advised to consider aspects pertaining to their specific datasets and hypotheses. As an example, use of GSR would remove signal of interest pertaining to some pharmacological and pathological states of altered consciousness (Luppi et al., 2019), (Luppi and Stamatakis, 2021; Tanabe et al., 2020)). Likewise, use of GSR has also reduced sensitivity to differences between Alzheimer’s disease patients and controls in terms of modularity and small worldness (Chen et al., 2018), and a recent study observed reduced generalisability of graph-theoretical properties across sites, sessions, and paradigms when GSR was used (22). A comprehensive evaluation of the relative advantages and drawbacks of GSR is beyond the scope of this paper, and the reader is referred to Fox & Murphy (2017) for a discussion. However, this issue highlights the broader point we raised above: test-retest reliability is only one of several aspects that should be taken into account before deciding whether to include a given step as part of one’s chosen pipeline for network analysis, and study-specific considerations will also come into play.

### Future directions

Future work may expand on the present results in several ways: for instance, by adopting multivariate connectivity estimators (Yoo et al., 2019) or methods from information decomposition capable of recovering different kinds of information sharing between regions (Mediano et al., 2019; Luppi et al., 2020) or the directionality of connections (transfer entropy, Granger causality, Dynamic Causal Modelling) (Friston et al., 2019) or disambiguating between direct and indirect connections (e.g. partial correlation; (Smith et al., 2011)). Finally, it remains to be determined how our results will generalise to the case of dynamic functional connectivity (Hutchison et al., 2013), and to frequency-specific networks obtained from EEG or MEG (Park et al., 2019),(Dimitriadis et al., 2018).

The generalizability of the proposed framework across populations is also worthy of future exploration. Compared to healthy controls, some clinical populations have demonstrated lower test-retest reliability, ranging from children with Attention Deficit Hyperactivity Disorder (Somandepalli et al., 2015) to individuals with Alzheimer’s disease (Conwell et al., 2018). Reliability across the lifespan should be also considered by comparing age-groups, as early evidence untangled age-related differences in test-retest reliability of rs-fMRI (Song et al., 2012). The choice of the optimal pipeline for test-retest reliability may therefore vary by clinical characteristics, which still remains to be ascertained using topology-based approaches such as portrait divergence. This is an important next step following the present work.

Overall, by enabling systematic evaluation of network processing steps in a way that does not require the arbitrary selection of specific network properties of interest, we hope that the topology-based framework proposed here will lead towards an objective consensus and more consistent practices in the network analysis of neuroimaging data.

## Conclusion

In conclusion, our study provides an exploratory framework searching for the best network construction pipeline across hundreds of candidates, with the aim of recovering a reliable brain network topology. Our findings indicate that no single choice can guarantee a reliable brain network topology, but the combination of several specific steps is the key to obtaining reliable brain networks. Finally, we revealed substantial differences across pipelines in terms of their ability to recover similar network topologies across different scans of the same individual - even within the same hour. This distinction of highly reliable versus poor pipelines further enhances the importance of identifying suitable network construction pipelines.

## Supporting information

Interactive Supplementary Tables

## Acknowledgements

This work was supported by the Gates Cambridge Trust [AIL]; the Canadian Institute for Advanced Research (CIFAR; grant RCZB/072 RG93193) [to DKM and EAS]; The National Institute for Health Research (NIHR, UK), Cambridge Biomedical Research Centre and NIHR Senior Investigator Awards [DKM]; The British Oxygen Professorship of the Royal College of Anaesthetists [DKM]; The Stephen Erskine Fellowship, Queens’ College, University of Cambridge [EAS]; the Medical Research Council Doctoral Training Grant (#RG86932) [HMG]; Pinsent Darwin Award [HMG]; MRC grant MR/K004360/1 (Behavioural and Neurophysiological Effects of Schizophrenia Risk Genes: A Multi-locus, Pathway Based Approach) [SID] and by a MARIE-CURIE COFUND EU-UK Research Fellowship [SID]. Acquisition of the NYU Test-Retest dataset was funded by Stavros S. Niarchos Foundation, the Leon Lowenstein Foundation, NARSAD (The Mental Health Research Association) grants to F.Xavier Castellanos; and Linda and Richard Schaps, Jill and Bob Smith, and the Taubman Foundation gifts to F.Xavier Castellanos.

## Author contributions

AIL, EAS, SID conceived the study. AIL, HMG, SID and EAS designed the methodology and the analysis. ARDP contributed to data analysis. AIL and HMG analysed the data. EAS, AEM participated in data collection. AIL, SID, HMG wrote the manuscript with feedback from all co-authors.

## Data and code availability

The NYU dataset is freely available from the International Neuroimaging Data-Sharing Initiative (INDI) (http://www.nitrc.org/projects/nyu_trt). The Cambridge dataset is available upon request from author EAS. The CONN toolbox is freely available online (http://www.nitrc.org/projects/conn). Python code for the portrait divergence is freely available online (https://github.com/bagrow/network-portrait-divergence). MATLAB code for the Orthogonal Minimum Spanning Tree thresholding is freely available online (https://github.com/stdimitr/topological_filtering_networks). The Brain Connectivity Toolbox code used for graph-theoretical analyses is freely available online (https://sites.google.com/site/bctnet/).

## References

Andellini, M., Cannatà, V., Gazzellini, S., Bernardi, B., Napolitano, A., 2015. Test-retest reliability of graph metrics of resting state MRI functional brain networks: A review. J. Neurosci. Methods 253, 183–192. doi:10.1016/j.jneumeth.2015.05.020

Arslan, S., Ktena, S.I., Makropoulos, A., Robinson, E.C., Rueckert, D., Parisot, S., 2018. Human brain mapping: A systematic comparison of parcellation methods for the human cerebral cortex. Neuroimage 170, 5–30. doi:10.1016/j.neuroimage.2017.04.014

Bagrow, J.P., Bollt, E.M., 2019. An information-theoretic, all-scales approach to comparing networks. Appl. Netw. Sci. 4, 45. doi:10.1007/s41109-019-0156-x

Bassett, D.S., Sporns, O., 2017. Network neuroscience. Nat. Neurosci. 20, 353–364. doi:10.1038/nn.4502

Behzadi, Y., Restom, K., Liau, J., Liu, T.T., 2007. A component based noise correction method (CompCor) for BOLD and perfusion based fMRI. Neuroimage 37, 90–101. doi:10.1016/j.neuroimage.2007.04.042

Biswal, B., Yetkin, F.Z., Haughton, V.M., Hyde, J.S., 1995. Functional connectivity in the motor cortex of resting human brain using echo-planar MRI. Magn. Reson. Med. 34, 537–541. doi:10.1002/mrm.1910340409

Botvinik-Nezer, R., Holzmeister, F., Camerer, C.F., Dreber, A., Huber, J., Johannesson, M., Kirchler, M., Iwanir, R., Mumford, J.A., Adcock, R.A., Avesani, P., Baczkowski, B.M., Bajracharya, A., Bakst, L., Ball, S., Barilari, M., Bault, N., Beaton, D., Beitner, J., Benoit, R.G., Schonberg, T., 2020. Variability in the analysis of a single neuroimaging dataset by many teams. Nature 582, 84–88. doi:10.1038/s41586-020-2314-9

Braun, U., Plichta, M.M., Esslinger, C., Sauer, C., Haddad, L., Grimm, O., Mier, D., Mohnke, S., Heinz, A., Erk, S., Walter, H., Seiferth, N., Kirsch, P., Meyer-Lindenberg, A., 2012. Test-retest reliability of resting-state connectivity network characteristics using fMRI and graph theoretical measures. Neuroimage 59, 1404–1412. doi:10.1016/j.neuroimage.2011.08.044

Bullmore, E., Sporns, O., 2009. Complex brain networks: Graph theoretical analysis of structural and functional systems. Nat. Rev. Neurosci. 10, 186–198. doi:10.1038/nrn2575

Burianová, H., Faizo, N.L., Gray, M., Hocking, J., Galloway, G., Reutens, D., 2017. Altered functional connectivity in mesial temporal lobe epilepsy. Epilepsy Res. 137, 45–52. doi:10.1016/j.eplepsyres.2017.09.001

Cammoun, L., Gigandet, X., Meskaldji, D., Thiran, J.P., Sporns, O., Do, K.Q., Maeder, P., Meuli, R., Hagmann, P., 2012. Mapping the human connectome at multiple scales with diffusion spectrum MRI. J. Neurosci. Methods 203, 386–397. doi:10.1016/j.jneumeth.2011.09.031

Cao, H., McEwen, S.C., Forsyth, J.K., Gee, D.G., Bearden, C.E., Addington, J., Goodyear, B., Cadenhead, K.S., Mirzakhanian, H., Cornblatt, B.A., Carrión, R.E., Mathalon, D.H., McGlashan, T.H., Perkins, D.O., Belger, A., Seidman, L.J., Thermenos, H., Tsuang, M.T., van Erp, T.G.M., Walker, E.F., Cannon, T.D., 2019. Toward Leveraging Human Connectomic Data in Large Consortia: Generalizability of fMRI-Based Brain Graphs Across Sites, Sessions, and Paradigms. Cereb. Cortex 29, 1263–1279. doi:10.1093/cercor/bhy032

Cao, H., Plichta, M.M., Schäfer, A., Haddad, L., Grimm, O., Schneider, M., Esslinger, C., Kirsch, P., Meyer-Lindenberg, A., Tost, H., 2014. Test-retest reliability of fMRI-based graph theoretical properties during working memory, emotion processing, and resting state. Neuroimage 84, 888–900. doi:10.1016/j.neuroimage.2013.09.013

Carp, J., 2012. On the plurality of (methodological) worlds: estimating the analytic flexibility of FMRI experiments. Front. Neurosci. 6, 149. doi:10.3389/fnins.2012.00149

Chen, X., Liao, X., Dai, Z., Lin, Q., Wang, Z., Li, K., He, Y., 2018. Topological analyses of functional connectomics: A crucial role of global signal removal, brain parcellation, and null models. Hum. Brain Mapp. 39, 4545–4564. doi:10.1002/hbm.24305

Conwell, K., von Reutern, B., Richter, N., Kukolja, J., Fink, G.R., Onur, O.A., 2018. Test-retest variability of resting-state networks in healthy aging and prodromal Alzheimer’s disease. Neuroimage Clin. 19, 948–962. doi:10.1016/j.nicl.2018.06.016

De Vico Fallani, F., Latora, V., Chavez, M., 2017. A topological criterion for filtering information in complex brain networks. PLoS Comput. Biol. 13, e1005305. doi:10.1371/journal.pcbi.1005305

Demertzi, A., Tagliazucchi, E., Dehaene, S., Deco, G., Barttfeld, P., Raimondo, F., Martial, C., Fernández-Espejo, D., Rohaut, B., Voss, H.U., Schiff, N.D., Owen, A.M., Laureys, S., Naccache, L., Sitt, J.D., 2019. Human consciousness is supported by dynamic complex patterns of brain signal coordination. Sci. Adv. 5, eaat7603. doi:10.1126/sciadv.aat7603

Dimitriadis, S.I., Antonakakis, M., Simos, P., Fletcher, J.M., Papanicolaou, A.C., 2017a. Data-Driven Topological Filtering Based on Orthogonal Minimal Spanning Trees: Application to Multigroup Magnetoencephalography Resting-State Connectivity. Brain Connect. 7, 661–670. doi:10.1089/brain.2017.0512

Dimitriadis, S.I., Drakesmith, M., Bells, S., Parker, G.D., Linden, D.E., Jones, D.K., 2017b. Improving the Reliability of Network Metrics in Structural Brain Networks by Integrating Different Network Weighting Strategies into a Single Graph. Front. Neurosci. 11, 694. doi:10.3389/fnins.2017.00694

Dimitriadis, S.I., Routley, B., Linden, D.E., Singh, K.D., 2018. Reliability of Static and Dynamic Network Metrics in the Resting-State: A MEG-Beamformed Connectivity Analysis. Front. Neurosci. 12, 506. doi:10.3389/fnins.2018.00506

Dimitriadis, S.I., Salis, C., Tarnanas, I., Linden, D.E., 2017c. Topological Filtering of Dynamic Functional Brain Networks Unfolds Informative Chronnectomics: A Novel Data-Driven Thresholding Scheme Based on Orthogonal Minimal Spanning Trees (OMSTs). Front. Neuroinformatics 11, 28. doi:10.3389/fninf.2017.00028

Du, H.-X., Liao, X.-H., Lin, Q.-X., Li, G.-S., Chi, Y.-Z., Liu, X., Yang, H.-Z., Wang, Y., Xia, M.-R., 2015. Test-retest reliability of graph metrics in high-resolution functional connectomics: a resting-state functional MRI study. CNS Neurosci. Ther. 21, 802–816. doi:10.1111/cns.12431

Eickhoff, S.B., Yeo, B.T.T., Genon, S., 2018. Imaging-based parcellations of the human brain. Nat. Rev. Neurosci. 19, 672–686. doi:10.1038/s41583-018-0071-7

Fan, S., Hansen, M.E.B., Lo, Y., Tishkoff, S.A., 2016. Going global by adapting local: A review of recent human adaptation. Science 354, 54–59. doi:10.1126/science.aaf5098

Fornito, A., Zalesky, A., Breakspear, M., 2015. The connectomics of brain disorders. Nat. Rev. Neurosci. https://doi.org/10.1038/nrn3901

Fox, M.D., Raichle, M.E., 2007. Spontaneous fluctuations in brain activity observed with functional magnetic resonance imaging. Nat. Rev. Neurosci. 8, 700–711. doi:10.1038/nrn2201

Friston, K.J., Preller, K.H., Mathys, C., Cagnan, H., Heinzle, J., Razi, A., Zeidman, P., 2019. Dynamic causal modelling revisited. Neuroimage 199, 730–744. doi:10.1016/j.neuroimage.2017.02.045

Glasser, M.F., Coalson, T.S., Robinson, E.C., Hacker, C.D., Harwell, J., Yacoub, E., Ugurbil, K., Andersson, J., Beckmann, C.F., Jenkinson, M., Smith, S.M., Van Essen, D.C., 2016. A multi-modal parcellation of human cerebral cortex. Nature 536, 171–178. doi:10.1038/nature18933

Greicius, M.D., Srivastava, G., Reiss, A.L., Menon, V., 2004. Default-mode network activity distinguishes Alzheimer’s disease from healthy aging: evidence from functional MRI. Proc Natl Acad Sci USA 101, 4637–4642. doi:10.1073/pnas.0308627101

Hallquist, M.N., Hillary, F.G., 2019. Graph theory approaches to functional network organization in brain disorders: A critique for a brave new small-world. Netw. Neurosci. 3, 1–26. doi:10.1162/netn_a_00054

Holiga, Š., Hipp, J.F., Chatham, C.H., Garces, P., Spooren, W., D’Ardhuy, X.L., Bertolino, A., Bouquet, C., Buitelaar, J.K., Bours, C., Rausch, A., Oldehinkel, M., Bouvard, M., Amestoy, A., Caralp, M., Gueguen, S., Ly-Le Moal, M., Houenou, J., Beckmann, C.F., Loth, E., Dukart, J., 2019. Patients with autism spectrum disorders display reproducible functional connectivity alterations. Sci. Transl. Med. 11. doi:10.1126/scitranslmed.aat9223

Huang, Z., Zhang, J., Wu, J., Mashour, G.A., Hudetz, A.G., 2020. Temporal circuit of macroscale dynamic brain activity supports human consciousness. Sci. Adv. 6, eaaz0087. doi:10.1126/sciadv.aaz0087

Hutchison, R.M., Womelsdorf, T., Allen, E.A., Bandettini, P.A., Calhoun, V.D., Corbetta, M., Della Penna, S., Duyn, J.H., Glover, G.H., Gonzalez-Castillo, J., Handwerker, D.A., Keilholz, S., Kiviniemi, V., Leopold, D.A., de Pasquale, F., Sporns, O., Walter, M., Chang, C., 2013. Dynamic functional connectivity: promise, issues, and interpretations. Neuroimage 80, 360–378. doi:10.1016/j.neuroimage.2013.05.079

Johnson, K.A., Sperling, R.A., Sepulcre, J., 2013. Functional connectivity in Alzheimer’s disease: measurement and meaning. Biol. Psychiatry 74, 318–319. doi:10.1016/j.biopsych.2013.07.010

Karbasforoushan, H., Woodward, N.D., 2012. Resting-state networks in schizophrenia. Curr. Top. Med. Chem. 12, 2404–2414. doi:10.2174/156802612805289863

Kiviniemi, V., Starck, T., Remes, J., Long, X., Nikkinen, J., Haapea, M., Veijola, J., Moilanen, I., Isohanni, M., Zang, Y.-F., Tervonen, O., 2009. Functional segmentation of the brain cortex using high model order group PICA. Hum. Brain Mapp. 30, 3865–3886. doi:10.1002/hbm.20813

Korhonen, O., Saarimäki, H., Glerean, E., Sams, M., Saramäki, J., 2017. Consistency of Regions of Interest as nodes of fMRI functional brain networks. Netw. Neurosci. 1, 254–274. https://doi.org/10.1162/netn_a_00013

Korhonen, O., Zanin, M., Papo, D., 2021. Principles and open questions in functional brain network reconstruction. Hum. Brain Mapp. https://doi.org/10.1002/hbm.25462

Laumann, T.O., Snyder, A.Z., Mitra, A., Gordon, E.M., Gratton, C., Adeyemo, B., Gilmore, A.W., Nelson, S.M., Berg, J.J., Greene, D.J., McCarthy, J.E., Tagliazucchi, E., Laufs, H., Schlaggar, B.L., Dosenbach, N.U.F., Petersen, S.E., 2017. On the Stability of BOLD fMRI Correlations. Cereb. Cortex 27, 4719–4732. doi:10.1093/cercor/bhw265

Luppi, A.I., Craig, M.M., Pappas, I., Finoia, P., Williams, G.B., Allanson, J., Pickard, J.D., Owen, A.M., Naci, L., Menon, D.K., Stamatakis, E.A., 2019. Consciousness-specific dynamic interactions of brain integration and functional diversity. Nat. Commun 10, 4616. doi:10.1038/s41467-019-12658-9

Luppi, A.I., Mediano, P.A., Rosas, F.E., Allanson, J., Carhart-Harris, R.L., Williams, G.B., Craig, M.M., Finoia, P., Owen, A.M., Naci, L., Menon, D.K., Bor, D., Stamatakis, E.A., 2020. A Synergistic Workspace for Human Consciousness Revealed by Integrated Information Decomposition. bioRxiv 2020.11.25.398081. https://doi.org/10.1101/2020.11.25.398081

Luppi, A.I., Stamatakis, E.A., 2021. Combining network topology and information theory to construct representative brain networks. Netw. Neurosci. 5, 96–124. doi:10.1162/netn_a_00170

Manktelow, A.E., Menon, D.K., Sahakian, B.J., Stamatakis, E.A., 2017. Working Memory after Traumatic Brain Injury: The Neural Basis of Improved Performance with Methylphenidate. Front. Behav. Neurosci. 11, 58. doi:10.3389/fnbeh.2017.00058

Mediano, P.A.M., Rosas, F., Carhart-Harris, R.L., Seth, A.K., Barrett, A.B., 2019. Beyond integrated information: A taxonomy of information dynamics phenomena. arXiv http://arxiv.org/abs/1909.02297

Messaritaki, E., Dimitriadis, S.I., Jones, D.K., 2019. Optimization of graph construction can significantly increase the power of structural brain network studies. Neuroimage 199, 495–511. doi:10.1016/j.neuroimage.2019.05.052

Messé, A., 2020. Parcellation influence on the connectivity‐based structure–function relationship in the human brain. Hum. Brain Mapp. 41, 1167–1180. https://doi.org/10.1002/hbm.24866

Murphy, K., Fox, M.D., 2017. Towards a consensus regarding global signal regression for resting state functional connectivity MRI. Neuroimage 154, 169–173. doi:10.1016/j.neuroimage.2016.11.052

Nichols, T.E., Das, S., Eickhoff, S.B., Evans, A.C., Glatard, T., Hanke, M., Kriegeskorte, N., Milham, M.P., Poldrack, R.A., Poline, J.-B., Proal, E., Thirion, B., Van Essen, D.C., White, T., Yeo, B.T.T., 2017. Best practices in data analysis and sharing in neuroimaging using MRI. Nat. Neurosci. 20, 299–303. doi:10.1038/nn.4500

Noble, S., Scheinost, D., Constable, R.T., 2019. A decade of test-retest reliability of functional connectivity: A systematic review and meta-analysis. Neuroimage 203, 116157. doi:10.1016/j.neuroimage.2019.116157

Park, H.-J., Friston, K., 2013. Structural and functional brain networks: from connections to cognition. Science 342, 1238411. doi:10.1126/science.1238411

Park, Y.-H., Cha, J., Bourakova, V., Lee, J.-M., 2019. Frequency specific contribution of intrinsic connectivity networks to the integration in brain networks. Sci. Rep. 9, 4072. doi:10.1038/s41598-019-40699-z

Parkes, L., Fulcher, B., Yücel, M., Fornito, A., 2018. An evaluation of the efficacy, reliability, and sensitivity of motion correction strategies for resting-state functional MRI. Neuroimage 171, 415–436. doi:10.1016/j.neuroimage.2017.12.073

Petersen, S.E., Sporns, O., 2015. Brain networks and cognitive architectures. Neuron 88, 207–219. doi:10.1016/j.neuron.2015.09.027

Petrella, J.R., 2011. Use of graph theory to evaluate brain networks: a clinical tool for a small world? Radiology 259, 317–320. doi:10.1148/radiol.11110380

Popovych, O.V., Jung, K., Manos, T., Diaz-Pier, S., Hoffstaedter, F., Schreiber, J., Yeo, B.T.T., Eickhoff, S.B., 2021. Inter-subject and inter-parcellation variability of resting-state whole-brain dynamical modeling. Neuroimage 118201. doi:10.1016/j.neuroimage.2021.118201

Ran, Q., Jamoulle, T., Schaeverbeke, J., Meersmans, K., Vandenberghe, R., Dupont, P., 2020. Reproducibility of graph measures at the subject level using resting-state fMRI. Brain Behav. 10, 2336–2351. doi:10.1002/brb3.1705

Romero-Garcia, R., Atienza, M., Clemmensen, L.H., Cantero, J.L., 2012. Effects of network resolution on topological properties of human neocortex. Neuroimage 59, 3522–3532. doi:10.1016/j.neuroimage.2011.10.086

Rubinov, M., Sporns, O., 2010. Complex network measures of brain connectivity: uses and interpretations. Neuroimage 52, 1059–1069. doi:10.1016/j.neuroimage.2009.10.003

Rubinov, M., Sporns, O., 2011. Weight-conserving characterization of complex functional brain networks. Neuroimage 56, 2068–2079. doi:10.1016/j.neuroimage.2011.03.069

Saad, Z.S., Gotts, S.J., Murphy, K., Chen, G., Jo, H.J., Martin, A., Cox, R.W., 2012. Trouble at rest: how correlation patterns and group differences become distorted after global signal regression. Brain Connect. 2, 25–32. doi:10.1089/brain.2012.0080

Schaefer, A., Kong, R., Gordon, E.M., Laumann, T.O., Zuo, X.-N., Holmes, A.J., Eickhoff, S.B., Yeo, B.T.T., 2018. Local-Global Parcellation of the Human Cerebral Cortex from Intrinsic Functional Connectivity MRI. Cereb. Cortex 28, 3095–3114. doi:10.1093/cercor/bhx179

Shehzad, Z., Kelly, A.M.C., Reiss, P.T., Gee, D.G., Gotimer, K., Uddin, L.Q., Lee, S.H., Margulies, D.S., Roy, A.K., Biswal, B.B., Petkova, E., Castellanos, F.X., Milham, M.P., 2009. The resting brain: unconstrained yet reliable. Cereb. Cortex 19, 2209–2229. doi:10.1093/cercor/bhn256

Shirer, W.R., Jiang, H., Price, C.M., Ng, B., Greicius, M.D., 2015. Optimization of rs-fMRI Pre-processing for Enhanced Signal-Noise Separation, Test-Retest Reliability, and Group Discrimination. Neuroimage 117, 67–79. doi:10.1016/j.neuroimage.2015.05.015

Smith, S.M., Miller, K.L., Salimi-Khorshidi, G., Webster, M., Beckmann, C.F., Nichols, T.E., Ramsey, J.D., Woolrich, M.W., 2011. Network modelling methods for FMRI. Neuroimage 54, 875–891. doi:10.1016/j.neuroimage.2010.08.063

Smith, S.M., Nichols, T.E., Vidaurre, D., Winkler, A.M., Behrens, T.E.J., Glasser, M.F., Ugurbil, K., Barch, D.M., Van Essen, D.C., Miller, K.L., 2015. A positive-negative mode of population covariation links brain connectivity, demographics and behavior. Nat. Neurosci. 18, 1565–1567. doi:10.1038/nn.4125

Smith, S.M., Vidaurre, D., Beckmann, C.F., Glasser, M.F., Jenkinson, M., Miller, K.L., Nichols, T.E., Robinson, E.C., Salimi-Khorshidi, G., Woolrich, M.W., Barch, D.M., Uğurbil, K., Van Essen, D.C., 2013. Functional connectomics from resting-state fMRI. Trends Cogn Sci (Regul Ed) 17, 666–682. doi:10.1016/j.tics.2013.09.016

Somandepalli, K., Kelly, C., Reiss, P.T., Zuo, X.-N., Craddock, R.C., Yan, C.-G., Petkova, E., Castellanos, F.X., Milham, M.P., Di Martino, A., 2015. Short-term test-retest reliability of resting state fMRI metrics in children with and without attention-deficit/hyperactivity disorder. Dev. Cogn. Neurosci. 15, 83–93. doi:10.1016/j.dcn.2015.08.003

Song, J., Desphande, A.S., Meier, T.B., Tudorascu, D.L., Vergun, S., Nair, V.A., Biswal, B.B., Meyerand, M.E., Birn, R.M., Bellec, P., Prabhakaran, V., 2012. Age-related differences in test-retest reliability in resting-state brain functional connectivity. PLoS ONE 7, e49847. doi:10.1371/journal.pone.0049847

Sporns, O., 2011. The human connectome: a complex network. Ann. N. Y. Acad. Sci. 1224, 109–125. doi:10.1111/j.1749-6632.2010.05888.x

Stam, C.J., 2014. Modern network science of neurological disorders. Nat. Rev. Neurosci. 15, 683–695. doi:10.1038/nrn3801

Tanabe, S., Huang, Z., Zhang, Jun, Chen, Y., Fogel, S., Doyon, J., Wu, J., Xu, J., Zhang, Jianfeng, Qin, P., Wu, X., Mao, Y., Mashour, G.A., Hudetz, A.G., Northoff, G., 2020. Altered Global Brain Signal during Physiologic, Pharmacologic, and Pathologic States of Unconsciousness in Humans and Rats. Anesthesiology 132, 1392–1406. doi:10.1097/ALN.0000000000003197

Termenon, M., Jaillard, A., Delon-Martin, C., Achard, S., 2016. Reliability of graph analysis of resting state fMRI using test-retest dataset from the Human Connectome Project. Neuroimage 142, 172–187. doi:10.1016/j.neuroimage.2016.05.062

Tian, Y., Margulies, D.S., Breakspear, M., Zalesky, A., 2020. Topographic organization of the human subcortex unveiled with functional connectivity gradients. Nat. Neurosci. 23, 1421–1432. doi:10.1038/s41593-020-00711-6

Turk, E., van den Heuvel, M.I., Benders, M.J., de Heus, R., Franx, A., Manning, J.H., Hect, J.L., Hernandez-Andrade, E., Hassan, S.S., Romero, R., Kahn, R.S., Thomason, M.E., van den Heuvel, M.P., 2019. Functional connectome of the fetal brain. J. Neurosci. 39, 9716–9724. doi:10.1523/JNEUROSCI.2891-18.2019

Tzourio-Mazoyer, N., Landeau, B., Papathanassiou, D., Crivello, F., Etard, O., Delcroix, N., Mazoyer, B., Joliot, M., 2002. Automated anatomical labeling of activations in SPM using a macroscopic anatomical parcellation of the MNI MRI single-subject brain. Neuroimage 15, 273–289. doi:10.1006/nimg.2001.0978

Van Dijk, K.R.A., Hedden, T., Venkataraman, A., Evans, K.C., Lazar, S.W., Buckner, R.L., 2010. Intrinsic functional connectivity as a tool for human connectomics: theory, properties, and optimization. J. Neurophysiol. 103, 297–321. doi:10.1152/jn.00783.2009

Váša, F., Bullmore, E.T., Patel, A.X., 2018. Probabilistic thresholding of functional connectomes: Application to schizophrenia. Neuroimage 172, 326–340. doi:10.1016/j.neuroimage.2017.12.043

Vatansever, D., Menon, D.K., Manktelow, A.E., Sahakian, B.J., Stamatakis, E.A., 2015. Default mode dynamics for global functional integration. J. Neurosci. 35, 15254–15262. doi:10.1523/JNEUROSCI.2135-15.2015

Wang, J., Ren, Y., Hu, X., Nguyen, V.T., Guo, L., Han, J., Guo, C.C., 2017. Test-retest reliability of functional connectivity networks during naturalistic fMRI paradigms. Hum. Brain Mapp. 38, 2226–2241. doi:10.1002/hbm.23517

Wang, J.-H., Zuo, X.-N., Gohel, S., Milham, M.P., Biswal, B.B., He, Y., 2011. Graph theoretical analysis of functional brain networks: test-retest evaluation on short- and long-term resting-state functional MRI data. PLoS ONE 6, e21976. doi:10.1371/journal.pone.0021976

Welton, T., Kent, D., Constantinescu, C.S., Auer, D.P., Dineen, R.A., 2015. Functionally relevant white matter degradation in multiple sclerosis: a tract-based spatial meta-analysis. Radiology 275, 89–96. doi:10.1148/radiol.14140925

Whitfield-Gabrieli, S., Nieto-Castanon, A., 2012. Conn: a functional connectivity toolbox for correlated and anticorrelated brain networks. Brain Connect. 2, 125–141. doi:10.1089/brain.2012.0073

Woo, C.-W., Chang, L.J., Lindquist, M.A., Wager, T.D., 2017. Building better biomarkers: brain models in translational neuroimaging. Nat. Neurosci. 20, 365–377. doi:10.1038/nn.4478

Yeh, F.-C., Panesar, S., Fernandes, D., Meola, A., Yoshino, M., Fernandez-Miranda, J.C., Vettel, J.M., Verstynen, T., 2018. Population-averaged atlas of the macroscale human structural connectome and its network topology. Neuroimage 178, 57–68. doi:10.1016/j.neuroimage.2018.05.027

Yoo, K., Rosenberg, M.D., Noble, S., Scheinost, D., Constable, R.T., Chun, M.M., 2019. Multivariate approaches improve the reliability and validity of functional connectivity and prediction of individual behaviors. Neuroimage 197, 212–223. doi:10.1016/j.neuroimage.2019.04.060

