## Supplementary Figures S1-S6 for "Searching for Consistent Brain Network Topologies Across the Garden of (Shortest) Forking Paths"

### Supplementary Material 1 - Portrait divergence as a function of dataset and preprocessing step

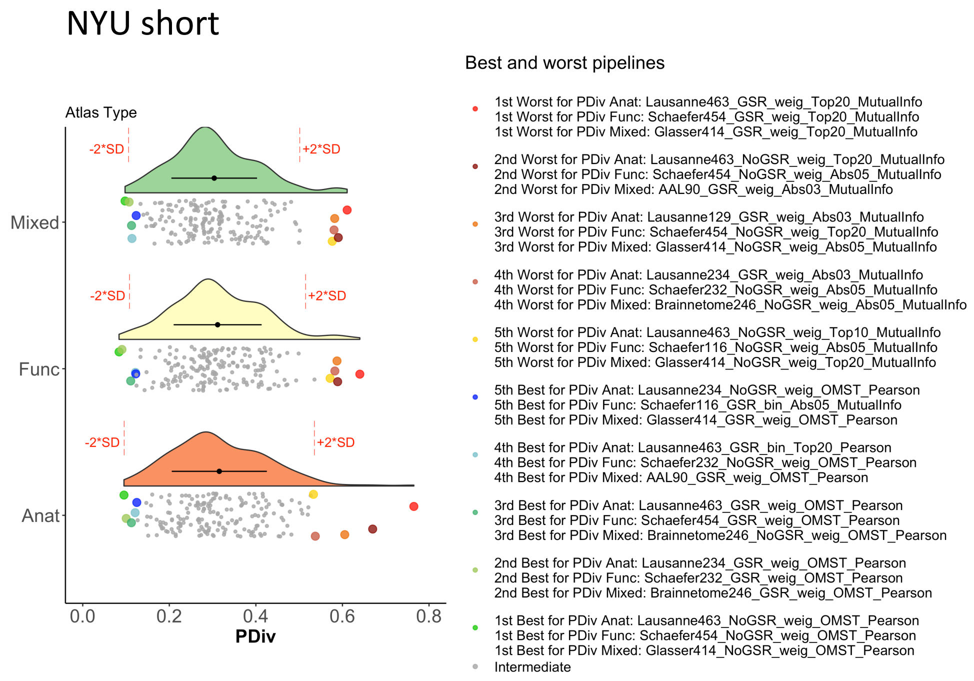


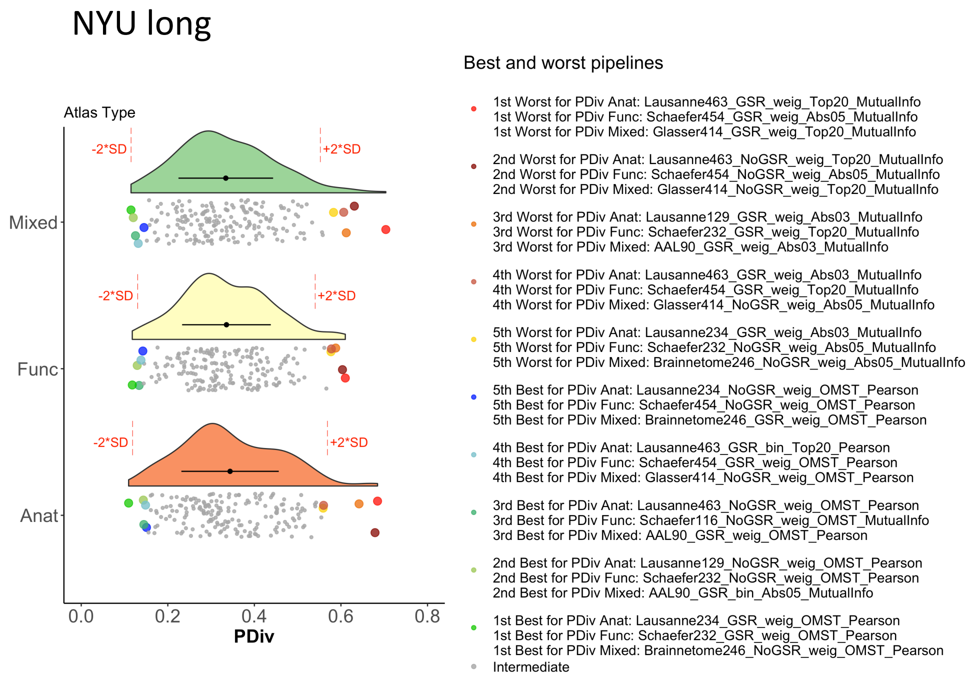


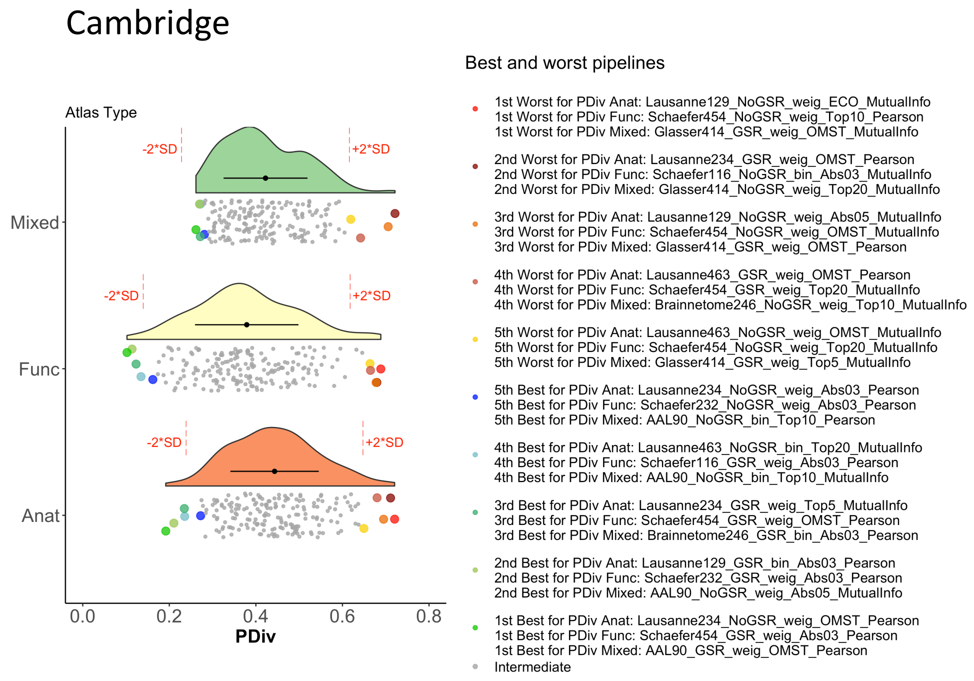


**Figure S1. Portrait divergence by atlas type and dataset.**

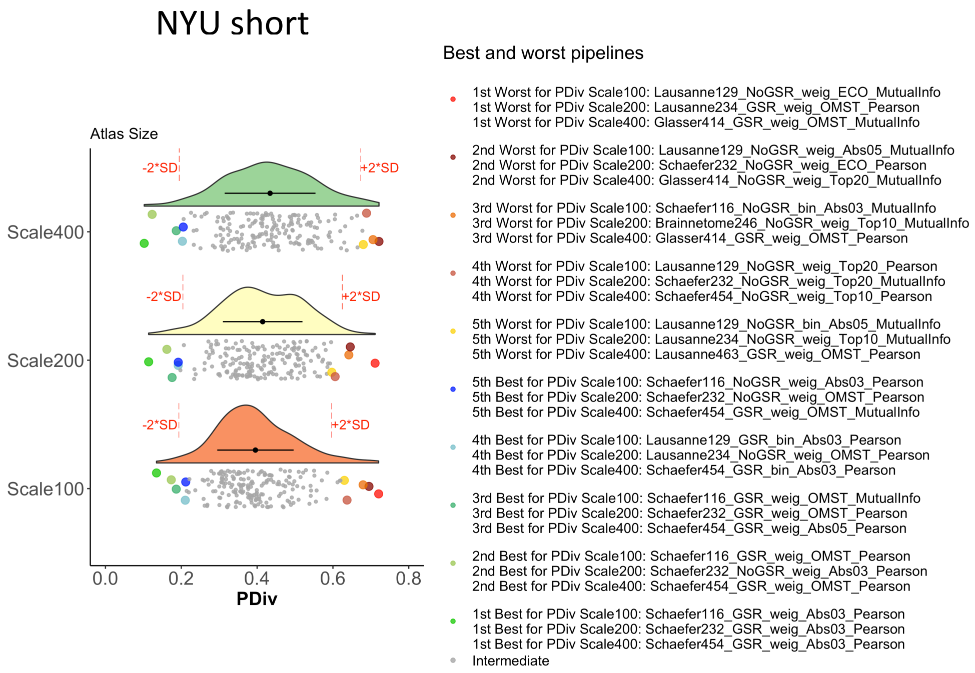


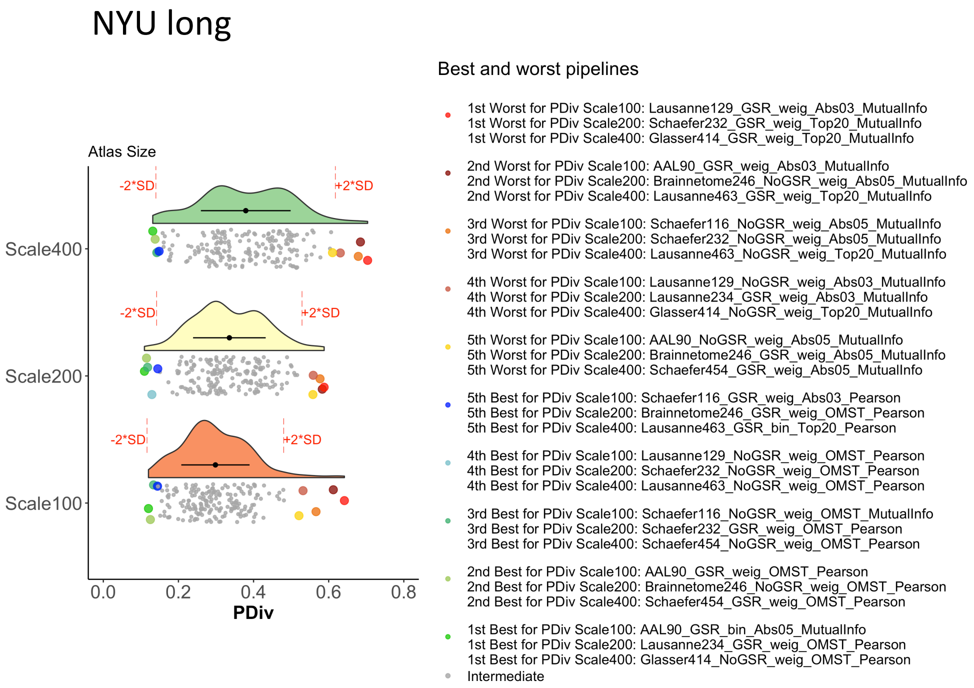


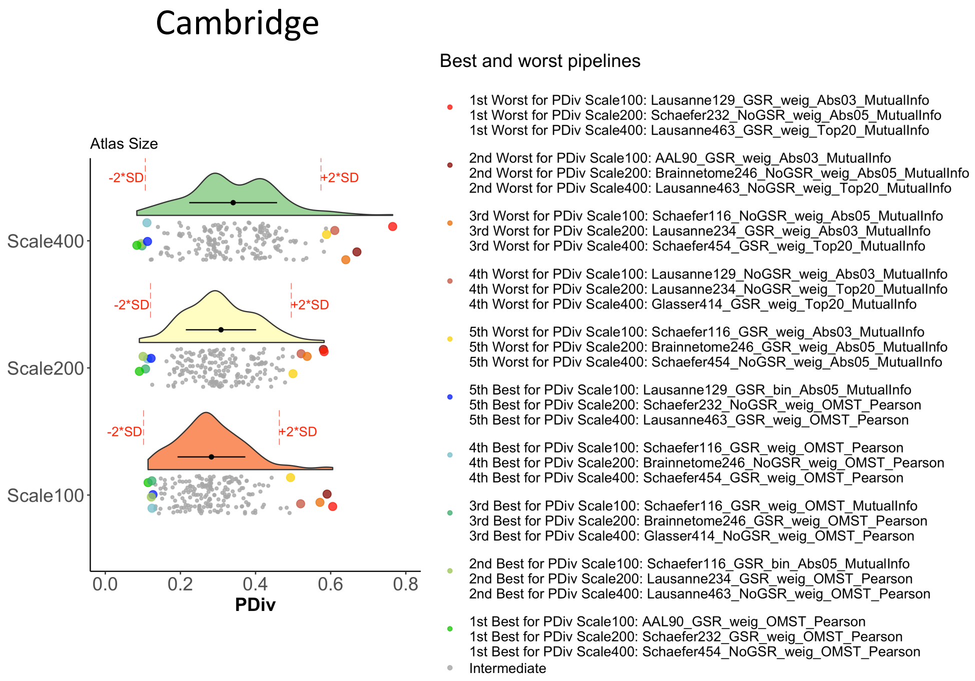


**Figure S2. Portrait divergence by atlas size and dataset.**

#### NYU short

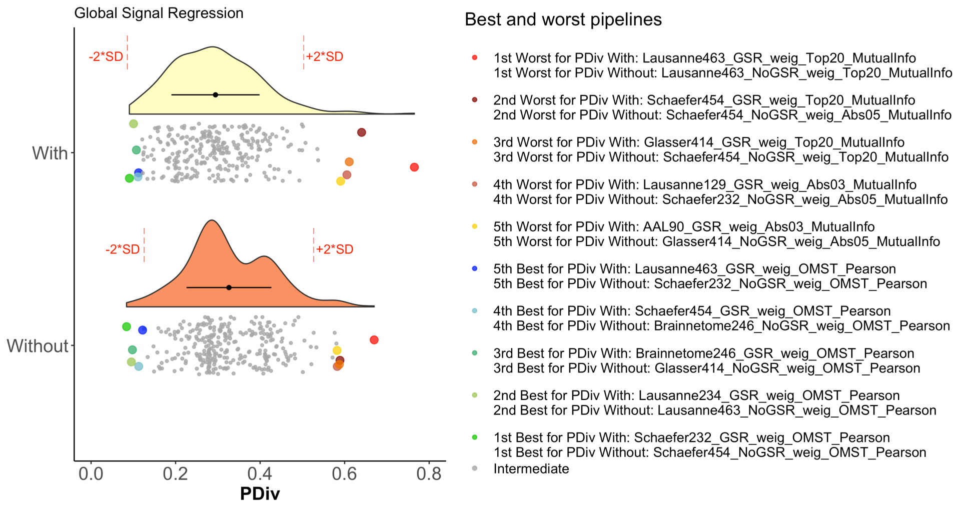


#### NYU long

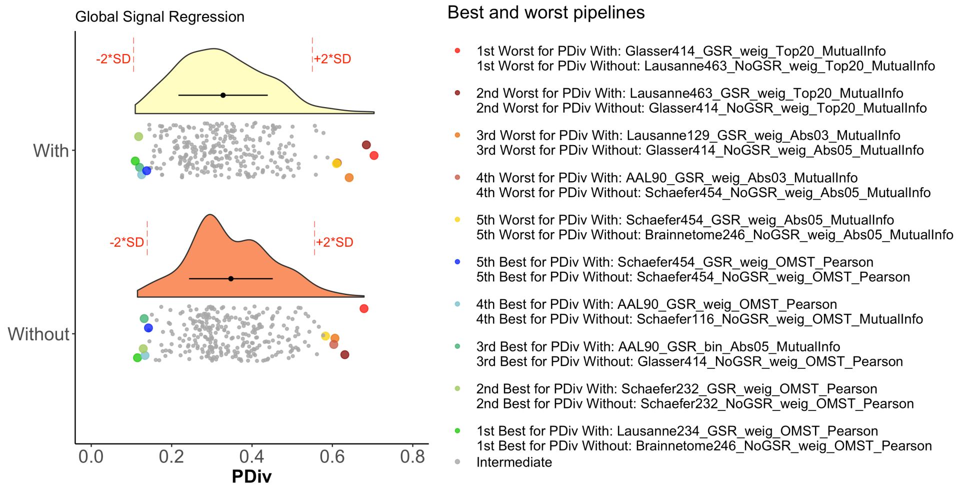


#### Cambridge

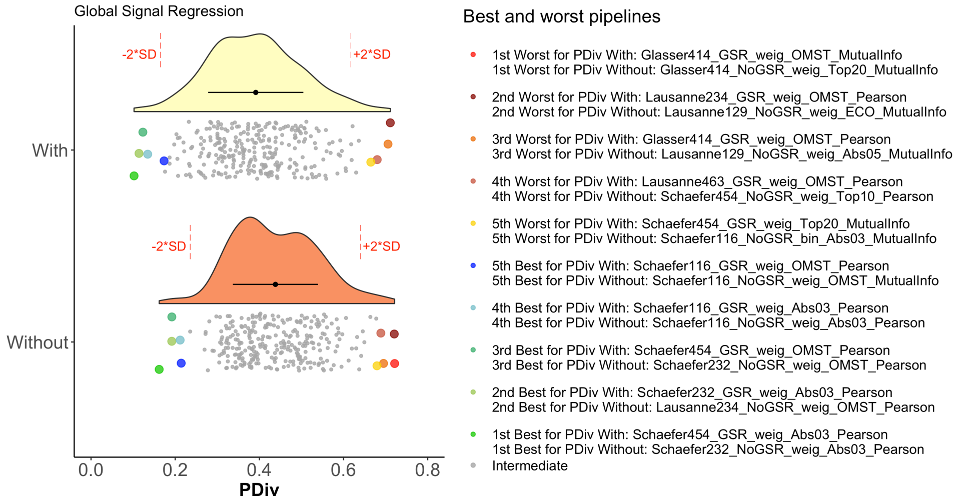


**Figure S3. Portrait divergence by GSR option and dataset.**

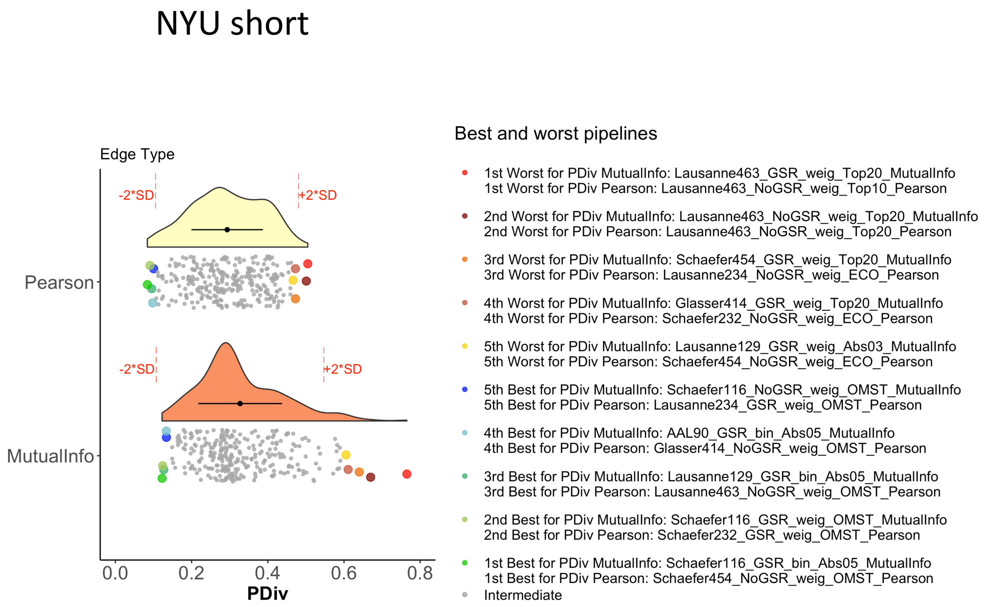


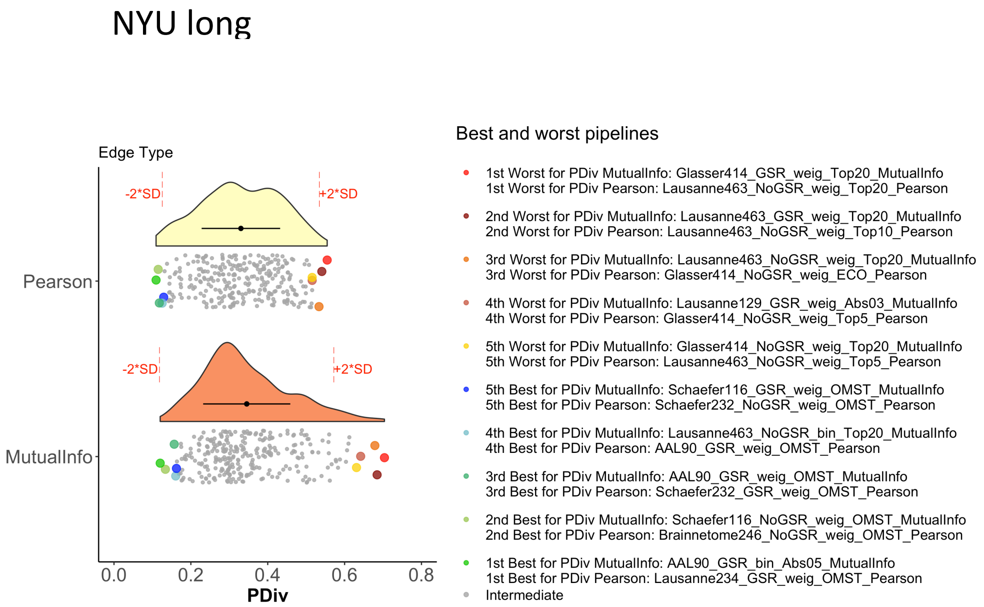


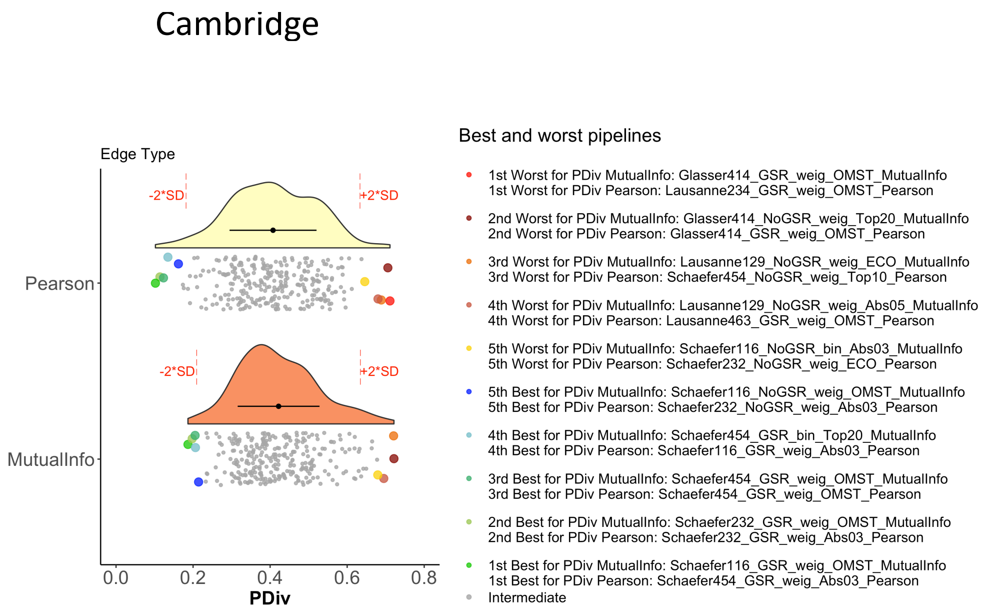


**Figure S4. Portrait divergence by edge type and dataset.**

#### NYU short

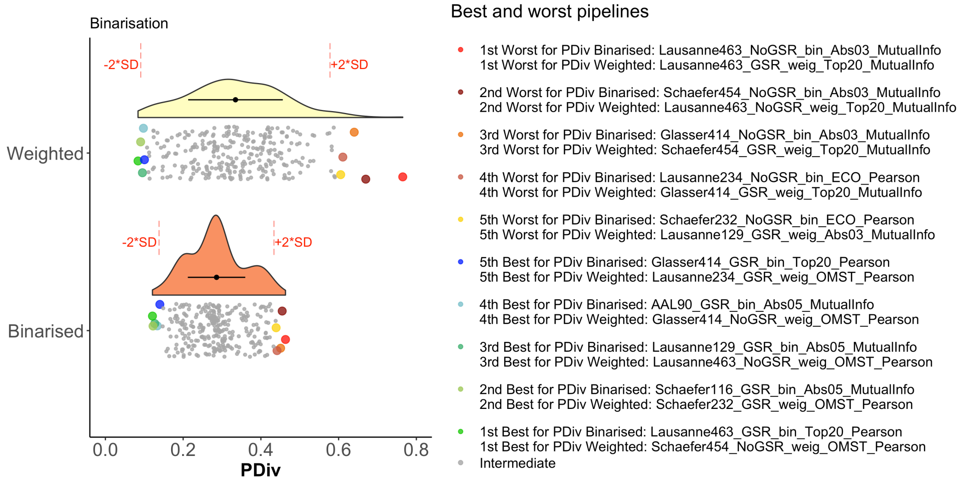


#### NYU long

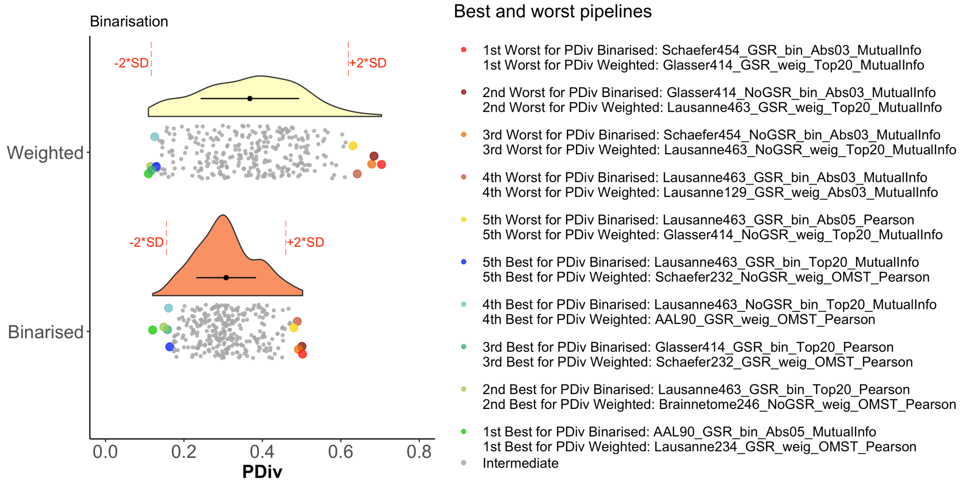


#### Cambridge

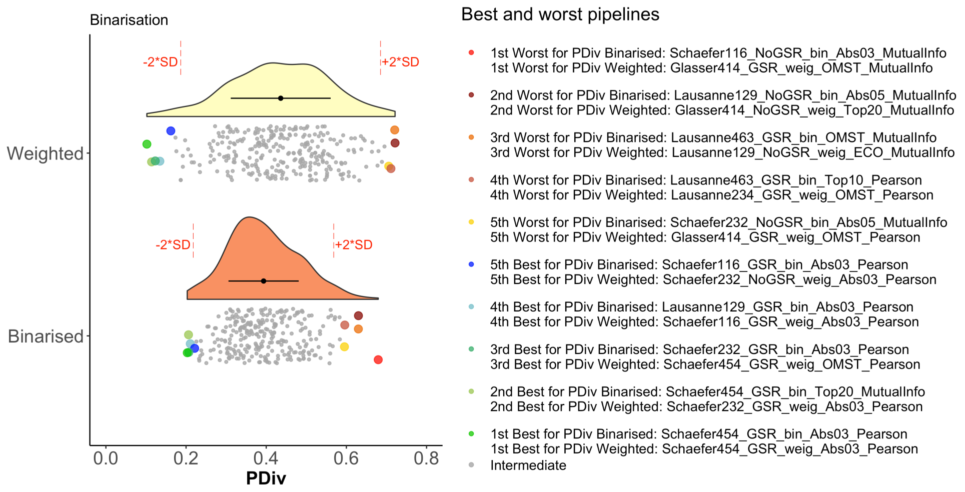


**Figure S5. Portrait divergence by binarization and dataset.**

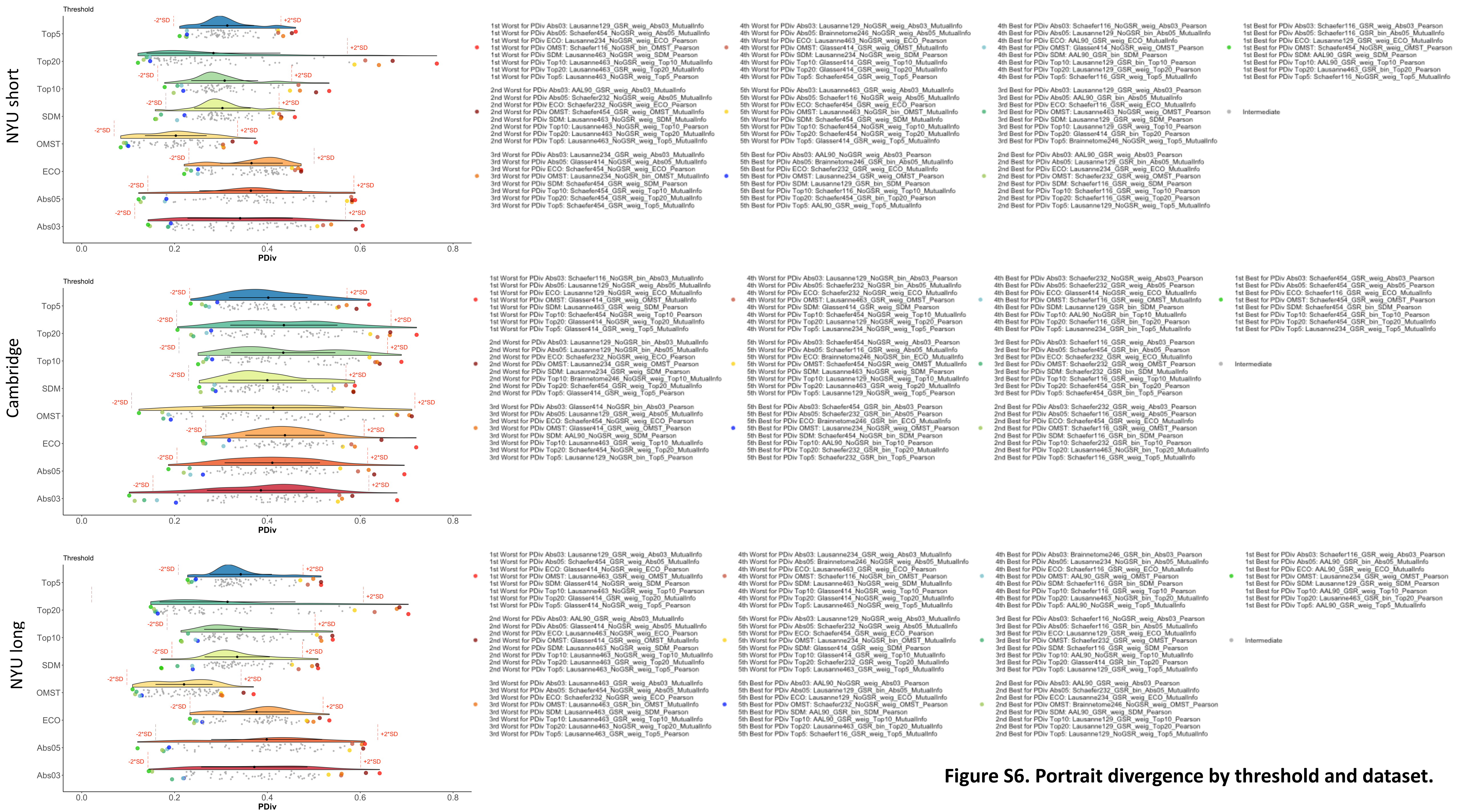


**Figure S6. Portrait divergence by threshold and dataset.**
